# Uncovering the mechanisms of clinically-relevant altered antibiotic responses of *Staphylococcus aureus* under wound infection-mimetic conditions

**DOI:** 10.64898/2025.12.22.696073

**Authors:** Colin D. Rieger, Ali Molaeitabari, Tanya E. S. Dahms, Omar M. El-Halfawy

## Abstract

Standard *in vitro* antimicrobial susceptibility testing (AST) using Mueller-Hinton broth (MHB) does not reflect infection-site conditions, and its results often do not correlate with therapeutic outcomes. Here, we compared the antibiotic susceptibility of methicillin-resistant *Staphylococcus aureus* (MRSA), a common chronic wound pathogen, in simulated wound fluid (SWF) resembling wound exudate versus MHB, revealing discordant AST results across six of nine tested antibiotic classes. The most significant were 128-fold increased resistance to tetracyclines and 256-fold sensitization to β-lactams in SWF. Tetracycline resistance was mediated by MntC, an extracellular manganese-binding protein, whereas β-lactam sensitization was driven by cell envelope remodelling in SWF. *Galleria mellonella* wound infection results matched the SWF susceptibility phenotypes, suggesting SWF better predicts *in vivo* wound infection therapeutic outcomes. These comprehensive phenotypic and mechanistic insights into MRSA antibiotic responses under wound-infection-mimetic conditions with direct *in vivo* validation identify a potential new antibiotic adjuvant target and may guide improved antibiotic therapy for MRSA wound infections.

## Introduction

Antibiotic resistance is rising at an alarming rate, and existing antibiotics are failing to treat infections ^1^. This crisis further complicates chronic infections, which pose significant healthcare and socioeconomic challenges ^2^. Chronic wounds are particularly troublesome because they have a prolonged inflammatory phase and delayed healing, leading to tissue fibrosis and non-healing ulcers ^3,4^. Wound exudates produced during healing are serum-based fluid rich in protein and growth factors, providing a suitable environment for bacterial colonization ^5^; such infections frequently fail to respond to treatment ^3^. Chronic wound infections are estimated to affect 1-2% of the world population at some point in their lives ^6^. Wound care poses a substantial economic burden on the healthcare system; wound care costs were estimated to be ∼$300B worldwide in 2019 ^7^ – such expenditures in the 10 highest spending countries alone increased from ∼$233B in 2019 to $301B in 2022 ^8^.

*Staphylococcus aureus,* a Gram-positive pathogen, is a leading cause of both hospital-and community-acquired infections worldwide, associated with bacteremia, endocarditis, skin and soft tissue infections (SSTIs), and other infections affecting almost all body sites ^9–11^. Notably, *S. aureus* is the leading cause of SSTIs, including complicated SSTIs such as chronic wounds and burn infections ^11–13^. If not treated, *S. aureus* wound infections can spread to bloodstream infections, which are often fatal ^14^. *S. aureus* developed multi-drug resistance, including to last-resort antibiotics, placing it on the World Health Organization’s list of ‘priority pathogens’ posing the greatest threat to human health ^15^. Methicillin-resistant *S. aureus* (MRSA, known for resistance to β-lactam antibiotics) is the leading cause of wound infections, posing severe clinical problems ^13^. Currently, the most commonly circulating community-associated MRSA lineage in North America is USA300 ^16^. The discovery of new antibiotics has lagged behind the constant rise in AMR prevalence ^10^. There is an urgent need for new antimicrobial solutions, including antibiotic adjuvants that revive currently failing antibiotics.

Bacterial antibiotic responses are often studied under standard *in vitro* conditions [e.g., Mueller-Hinton broth (MHB), composed of beef extract, casein hydrolysate, and starch^17^] that do not adequately reflect the infection site. These assays, standardized by agencies such as CLSI^18^, are routinely used in clinical labs worldwide to determine whether a clinical isolate is sensitive or resistant to an antibiotic, guiding antibiotic therapy. Nonetheless, the clinical outcome often does not correlate with standard antibiotic susceptibility test (AST) results, with reports of unexpected antibiotic therapeutic failure ^19–21^. In contrast, unexpected potency in patients despite AST-predicted resistance is far less frequently reported, as such antibiotics would typically be contraindicated. However, anecdotal reports describe instances in which an antibiotic empirically prescribed while waiting (up to 72 h) for AST results shows efficacy in patients before AST identifies the causative isolate as resistant to the same antibiotic. Together, standard AST erroneous predictions can lead to therapeutic failure or sideline much-needed *in vivo*-effective antibiotics.

Culture conditions and media composition dictate gene expression and dispensability profiles in bacteria to meet their survival requirements ^22,23^, thereby potentially altering bacterial antibiotic responses. Infection-mimetic media, such as simulated wound fluid (SWF) ^24^, have been developed, providing test conditions that are more clinically relevant than MHB for assessing bacterial responses. SWF is composed of 50% v/v fetal bovine serum in maximal recovery diluent, which is an isotonic peptone saline solution ^24^. Such a formulation of SWF mimics chronic wound exudates, as the latter are derived from serum and contain electrolytes at concentrations equivalent to those in serum, approximately half the total protein content of serum, and higher levels of growth factors for tissue regeneration ^25–29^. The inclusion of fetal bovine serum in this recipe provides additional components not present in MHB, such as growth factors, transport proteins, enzymes, hormones, fatty acids, and lipids, at biologically relevant levels ^30,31^.

Here, we used SWF to evaluate the antimicrobial susceptibility profile of MRSA under wound infection-mimetic conditions. We observed discordance in AST results for representative antibiotics across six of nine antibiotic classes tested in SWF compared to MHB against the MRSA USA300. The antibiotic classes showing the most significant discordant AST results included tetracyclines, to which USA300 was more resistant (up to a 128-fold shift), and β-lactams, to which the MRSA was unexpectedly more sensitive (up to a 256-fold shift), when tested in SWF. Interestingly, these antibiotic susceptibility shifts crossed the CLSI clinical breakpoint for representatives of both classes, suggesting they may result in altered therapeutic outcomes. Genome-wide assays and subsequent mechanistic characterizations revealed that the increased resistance to tetracyclines is mediated by MntC, an extracellular binding protein of a manganese ABC transporter system, whereas the drastic sensitization to β-lactams was driven by cell envelope remodelling in SWF. Further, we used a *Galleria mellonella* incision wound infection model, which shows wound healing events similar to those in animals ^32^ and recapitulates the hallmarks of trauma and infection observed in mammalian models ^33^, to characterize the observed *in vitro* phenotypes. Doxycycline, a tetracycline antibiotic, could not rescue wild-type USA300-infected Galleria larvae but significantly increased the survival of those infected with the *mntC*::Tn mutant, whereas the β-lactam cefuroxime was effective at rescuing a significant number of USA300-infected larvae. These *in vivo* results matched the *in vitro* phenotypes in SWF but not MHB, suggesting that this host-mimetic medium is a better predictor of *in vivo* MRSA wound infection therapeutic outcomes.

Together, this work addresses the relatively long-recognized shortcomings of the MHB-based standard AST by uncovering alterations in MRSA antibiotic susceptibility in wound exudate-like fluid relative to MHB. Further, this study not only describes the phenotypic differences between SWF and MHB but, more importantly, also systematically reveals the specific molecular mechanisms underlying the most significant discordant AST results between standard and infection-mimetic conditions at the whole-genome level, and directly links the uncovered *in vitro* phenotypes and mechanisms to *in vivo* efficacy. Therefore, our results provide a solid theoretical basis for improved antibiotic selection in the clinic and a potential new target for adjuvant therapy against drug-resistant MRSA wound infections.

## Results

### Alterations in MRSA sensitivity to antibiotics in SWF

We first compared AST results of representative antibiotics from nine classes against USA300 cultured in MHB and SWF (Table 1). Six of the nine classes showed discordance in minimum inhibitory concentration (MIC) determined in the two media (MIC ≥ 4-fold difference between SWF and MHB, Table 1). These MIC discordances included a 128-, 16-, and 4-fold sensitization in SWF to azithromycin (a macrolide), colistin (a polymyxin antimicrobial peptide), and daptomycin (a lipopeptide), respectively, and an 8-fold increase in rifampicin MIC in SWF against USA300 (Table 1). These MIC shifts did not cross the CLSI breakpoints for the four antibiotic classes in USA300 (Table 1). Yet, similar MIC fold shifts in other *S. aureus* backgrounds that are less susceptible to these antibiotics may alter the predicted antibiotic susceptibility, warranting future testing in other *S. aureus* lineages. The remaining two antibiotic classes showing the most significant fold shift in MIC in SWF relative to MHB are tetracyclines that showed an increase in MIC when tested in SWF (e.g., a 128-fold shift for tetracycline from 0.25 μg/mL in MHB to 32 μg/mL in SWF) and β-lactams that showed a decrease in MIC in SWF (e.g., a 256-fold shift for cefuroxime from 1024 μg/mL in MHB to 4 μg/mL in SWF, Table 1). Importantly, the MIC fold shifts crossed the CLSI clinical breakpoint for representatives of these two classes (Table 1). Together, we chose to focus on these two classes to uncover the mechanism underlying the observed altered antibiotic susceptibility under the wound-mimetic condition.

**Table 1.** USA300 MIC assays conducted in SWF relative to those in MHB MICs from at least three independent experiments.

| Antibiotic class | Antibiotic | MIC, µg/mL |  | Fold shift* | CLSI breakpoint, µg/mL |  |  |
| --- | --- | --- | --- | --- | --- | --- | --- |
|  |  | MHB | SWF |  | R, ≥ | S, ≤ | Ref |
| Lipopeptides | Daptomycin | 4 | 1 | 4 ↓ | - | 1 | <sup>91</sup> |
| Fluoroquinolones | Ciprofloxacin | 32 | 16 | 2 ↓ | 4 | 1 | <sup>91</sup> |
| Glycopeptides | Vancomycin | 2 | 1 | 2 ↓ | 16 | 2 | <sup>91</sup> |
| Rifampins | Rifampicin | 0.008 | 0.064 | 8 ↑ | 4 | 1 | <sup>91</sup> |
| β-lactams | Cefuroxime | 1024 | 4 | 256 ↓ | 32 | 8 | <sup>92</sup> |
|  | Oxacillin | 64 | 1 | 64 ↓ | 4 | 2 | <sup>91</sup> |
| Polymyxins | Colistin | 512 | 32 | 16 ↓ |  |  |  |
| Tetracyclines | Tetracycline | 0.25 | 32 | 128 ↑ | 16 | 4 | <sup>91</sup> |
|  | Doxycycline | 0.125 | 4 | 32 ↑ | 16 | 4 | <sup>91</sup> |
|  | Omadacycline | 0.25 | 0.5 | 2 ↑ |  |  |  |
| Macrolides | Azithromycin | 2 | 0.0156 | 128 ↓ | 8 | 2 | <sup>91</sup> |
| Aminoglycosides | Kanamycin | 4 | 4 | No shift | 64 | 16 | <sup>18</sup> |

\* Fold increase or decrease in SWF relative to MHB.

Tetracycline antibiotics inhibit protein synthesis by binding to the 30S ribosome subunit ^34^. Doxycycline is a second-generation tetracycline that showed a 32-fold increase in MIC against USA300 from 0.125 μg/mL in MHB to 4 μg/mL in SWF (Table 1). In addition, we assessed the third-generation tetracycline, omadacycline, and found that its MIC against USA300 doubled in SWF relative to MHB (Table 1). These results demonstrate that the increased resistance in SWF is conserved across the three generations of tetracyclines. Doxycycline is more clinically relevant than tetracycline for treating MRSA skin and soft tissue infections (SSTIs) ^35^; therefore, we used it as a representative of this class in the remainder of this study. On the other hand, cefuroxime is a β-lactam antibiotic that binds to penicillin-binding protein 2, inhibiting bacterial cell wall synthesis ^36^. Oxacillin, another β-lactam, is used by CLSI to distinguish MRSA from methicillin-sensitive *S. aureus* (MSSA) ^18^; it showed a 64-fold decrease in MIC against USA300 in SWF (from 64 μg/mL in MHB to 1 μg/mL in SWF), bringing the MIC below the CLSI MRSA breakpoint (Table 1). We initially tested both β-lactams and then focused on cefuroxime for subsequent mechanistic testing.

To determine how conserved the doxycycline and β-lactam susceptibility alterations in SWF are across multiple MRSA strains, we determined the MICs of antibiotics against a panel of isolates representing the most commonly circulating lineages of MRSA in Canada [CMRSA strains ^37^] in both MHB and SWF. All CMRSA strains showed increased resistance (≥ 8-fold shift in MIC) to doxycycline when tested in SWF relative to MHB (Table 2). All CMRSA strains exhibited increased sensitivity to both β-lactams in SWF, except for CMRSA 1 and CMRSA 4, with MIC shifts in CMRSA 5, 7, 8, and 10 crossing the CLSI breakpoint for one or both antibiotics (Table 2). Given the magnitude of antibiotic susceptibility alterations conserved across strains from multiple MRSA lineages, we next sought to determine the mechanisms of doxycycline and cefuroxime susceptibility alterations in the wound-mimetic medium in MRSA using USA300 as a representative being one of the most common community-associated MRSA strains.

**Table 2.** MIC of cefuroxime, oxacillin, and doxycycline against isolates representing the most commonly circulating lineages of MRSA in Canada. MICs average of three independent experiments.

| Strain | Lineage | MIC in MHB (µg/mL) |  |  | MIC in SWF (µg/mL) |  |  |
| --- | --- | --- | --- | --- | --- | --- | --- |
|  |  | cefuroxime | oxacillin | doxycycline | cefuroxime | oxacillin | doxycycline |
| CMRSA 1 | USA600 | >2048 | 1024 | 0.25 | >2048 | 1024 | 2 |
| CMRSA 2 | USA100/800 | >2048 | 512 | 0.25 | 16 | 32 | 4 |
| CMRSA 3 |  | >2048 | 512 | 16 | 512 | 256 | >128 |
| CMRSA 4 | USA200 | 2048 | 256 | 0.25 | 2048 | 256 | 4 |
| CMRSA 5 | USA500 | 256 | 32 | 8 | 8 | 1 | >128 |
| CMRSA 6 |  | >2048 | 512 | 16 | 1024 | 256 | >128 |
| CMRSA 7 | USA400/MW2 | 1024 | 64 | 0.25 | 8 | 2 | 2 |
| CMRSA 8 | EMRSA15 | 1024 | 64 | 0.25 | 4 | 8 | 2 |
| CMRSA 9 |  | 2048 | 256 | 0.25 | 16 | 32 | 4 |
| CMRSA 10 | USA300 | 1024 | 64 | 0.25 | 4 | 2 | 4 |

### A chemogenomic screen to uncover the mechanism of increased resistance to doxycycline in SWF

To investigate the increased resistance of USA300 to doxycycline when cultured in SWF, we screened the Nebraska Transposon Mutant Library (NTML), a sequence-defined transposon insertion library comprising 1920 mutants that cover the non-essential genome of USA300 ^38^, at supra-inhibitory (in MHB) and sub-inhibitory (in SWF) concentrations of doxycycline. The NTML supra-inhibitory screen in MHB (at 1 µg/mL doxycycline, 8X the MIC in MHB), which aimed to identify putative sensitivity determinants, and follow-up dose-response assays did not reveal any mutants more resistant to doxycycline in MHB relative to the wild-type (Fig. S1A-C). On the other hand, the NTML sub-inhibitory screen in SWF (at 0.125 µg/mL doxycycline, 1/32^nd^ the MIC in SWF), which aimed to identify putative determinants of doxycycline resistance, and follow-up MIC assays revealed that *mntB*::Tn and *mntC*::Tn were sensitive to doxycycline in SWF relative to USA300 (Fig. 1A-C and Fig. S2). To verify the observed phenotypes, we genetically complemented *mntB* and *mntC* in their transposon mutants. While *mntC* complementation restored doxycycline resistance of *mntC*::Tn, *mntB* alone was not sufficient to restore doxycycline resistance in *mntB*::Tn, but rather both *mntB* and *mntC* were required (Fig. S3A-C). As such, the *mntB*::Tn phenotype may be due to polar effects from the transposon insertion into *mntB,* disrupting the downstream *mntC* (Fig. S3C). Together, this work identifies MntC as the determinant of increased doxycycline resistance in the wound-like medium.

**Fig. 1.**
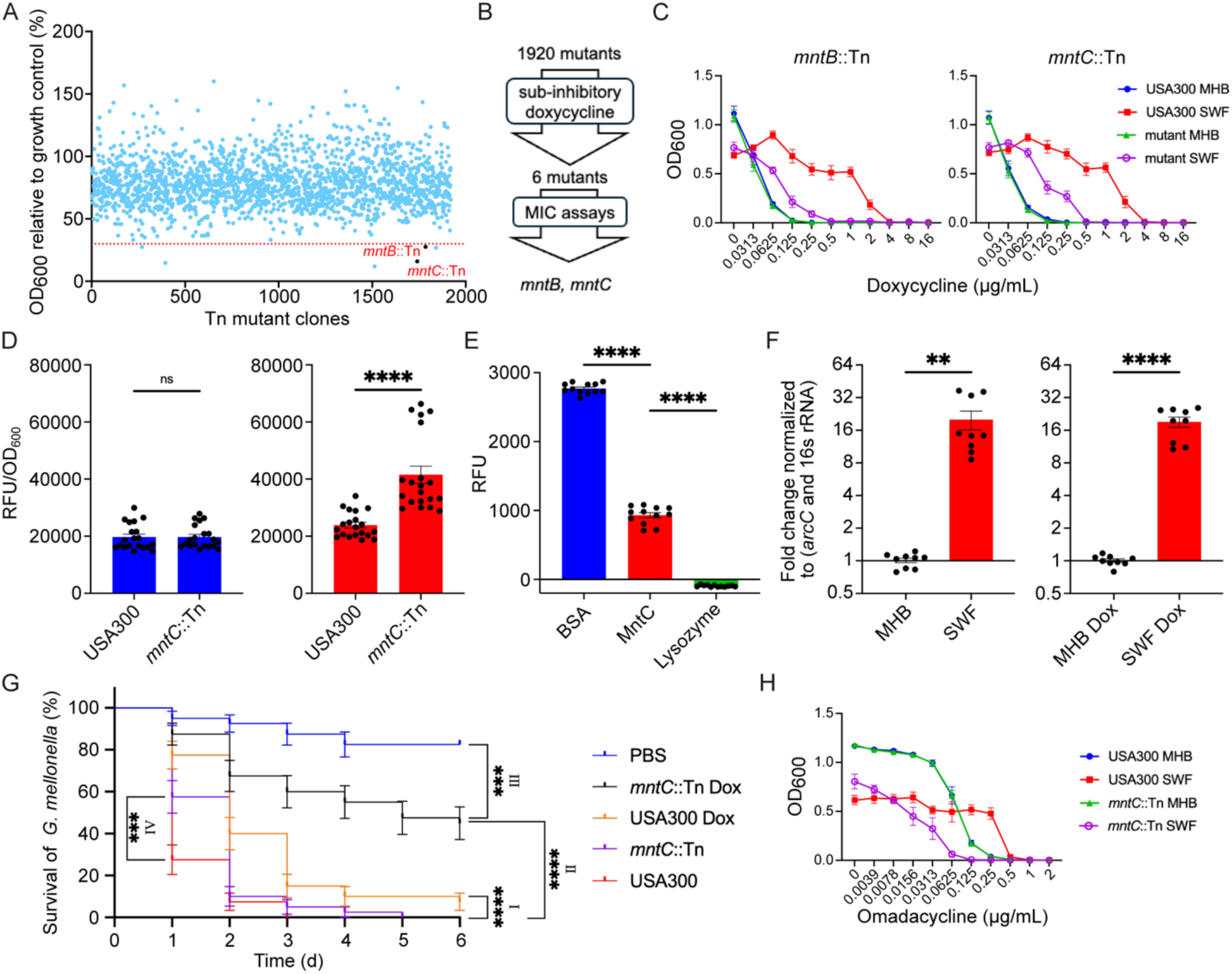
MntC increases USA300 resistance to doxycycline in SWF. A) A chemogenomic screen of the Nebraska Transposon Mutant Library with doxycycline at 1/32^th^ the MIC in SWF. B) Workflow of the chemogenomic screen and follow-up assays that lead to the putative determinants of doxycycline resistance. C) MIC assays of determinants of USA300 doxycycline resistance in SWF. n = 9 from three independent experiments, shown as mean ± SEM. D) Intracellular fluorescence of doxycycline relative fluorescent units (RLU)/OD_600_ in USA300 and *mntC*::Tn cultured in MHB (blue) or SWF (red). n = 20 from 4 independent experiments. E) Doxycycline binding assay in HEPES buffer showing increased background-subtracted RFU upon doxycycline binding with the ligands bovine serum albumin (positive control), MntC, and lysozyme (negative control). n = 12. F) Fold change in *mntC* transcription from USA300 in SWF and MHB with or without doxycycline (relative to averaged signals in MHB). n = 9 from three independent experiments. G) Kaplan-Meier survival curve of *G. mellonella* treated with doxycycline or vehicle control, shown as mean ± SE. n = 40 from 4 independent experiments. H) MIC assays of omadacycline against USA300 and *mntC*::Tn cultured in SWF and MHB. n = 6 from three independent experiments, shown as mean ± SEM. Significant difference were determined between USA300 cultured in MHB and SWF for D, E, F by Welch’s t-test, and in G (I; USA300 and USA300 Dox), (II; *mntC*::Tn and *mntC*::Tn Dox) and (III; *mntC*::Tn Dox and PBS) by Logrank (Mantel-Cox) test and (IV; *mntC*::Tn and USA300) at day 1 by two-way ANOVA and the Tukey post hoc test. p <0.0001 (****), p <0.001 (***), p <0.01 (**), p <0.05 (*).

### MntC increases doxycycline resistance in wound-like medium

Next, we set out to determine the mechanism by which MntC increases doxycycline resistance in SWF. MntABC, an ABC transporter that imports manganese, is comprised of an ATP-binding protein (MntA), a transmembrane protein (MntB), and an extracellular binding protein (MntC) ^39^; therefore, we initially hypothesized that manganese or other divalent cations imported by MntC contributed to the increased resistance to doxycycline. However, exogenously supplementing manganese, calcium, magnesium, cobalt, iron, copper, and zinc, in dose-dependent checkerboard assays with doxycycline against wild-type USA300 and *mntC*::Tn in MHB did not alter the susceptibility to doxycycline except for iron, which showed antagonism in both strains suggesting the observed iron effect is MntC-independent (Fig. S4). Conversely, higher concentrations of manganese antagonized doxycycline in SWF both in the wild-type and the *mntC* mutant (FICI = 6.75 and 32.5, respectively; Fig. S5). Such effect might be due to compensation of the sequestration of manganese in SWF due to chelators, such as calprotectin, likely present in fetal bovine serum. Together, manganese and other divalent cations appear to play no role in the MntC-mediated doxycycline resistance observed in SWF.

Next, we hypothesized that MntC sequesters doxycycline, lowering its intracellular concentration, thereby increasing resistance to that antibiotic in bacteria cultured in SWF. To address this hypothesis, we first determined the intracellular concentration of doxycycline in *mntC*::Tn relative to the wild-type cultured in MHB and SWF. In SWF, *mntC*::Tn had a greater intracellular concentration of doxycycline relative to USA300, whereas both the wild-type and mutant strains showed similar intracellular doxycycline content in MHB (Fig. 1D), aligning with our previous results showing a SWF-specific role of MntC in doxycycline resistance and supporting our hypothesis of a potential sequestration of doxycycline by MntC. Next, we assessed the ability of purified recombinant MntC to bind doxycycline using a fluorescent binding assay conducted in HEPES buffer in which doxycycline fluorescence increased upon binding to a ligand. Using bovine serum albumin and lysozyme as positive ^40^ and negative controls, respectively, in this assay, we revealed that MntC binds doxycycline (Fig. 1E), which likely reduces its intracellular concentration (Fig. 1D). Given that such a binding interaction does not require any component of SWF, as it occurred in HEPES buffer, we posited that MntC-mediated increased doxycycline resistance occurs in SWF only due to an upregulation of *mntC* under wound-like conditions. Hence, we assessed the transcriptional differences in *mntC* using qRT-PCR. We revealed that *mntC* was upregulated by ∼20-fold in wild-type USA300 cultured in SWF relative to MHB, irrespective of doxycycline treatment (Fig. 1F), demonstrating that the *mntC* upregulation is induced by the wound-like culture condition rather than a specific response to doxycycline. In fact, doxycycline exposure downregulated *mntC* in USA300 in both SWF and MHB to a similar extent (∼4-fold; Fig. S6). In summary, the increase in doxycycline resistance in the wound-like medium is due to an upregulation of *mntC*, whereby the overexpressed MntC can sequester doxycycline, decreasing its effective intracellular concentration.

### MntC contributes to doxycycline resistance *in vivo* in a *Galleria mellonella* wound infection model

Next, we sought to assess whether MntC contributed to doxycycline resistance *in vivo* in a wound infection model. To that end, we tested the *mntC*::Tn mutant in a *Galleria mellonella* incision wound infection model. Galleria mounts both cellular and humoral innate immune responses closely resembling those of vertebrates and thrives at mammalian body temperature ^41,42^. Incision wounds have been histologically characterized in Galleria larvae, showing wound healing events similar to those in animals ^32^, and their wound model recapitulates the hallmarks of trauma and infection observed in mammalian models ^33^. Infection with either the wild-type USA300 or the *mntC*::Tn mutant, both treated with the vehicle control (Glaxal^TM^ Base non-medicated cream), led to 100% larval fatality by the end of the experiment (at 6 days); however, we observed a significant increase in survival of larvae infected with *mntC*::Tn relative to USA300 at day 1 (Fig. 1G), suggesting a role of MntC in virulence. Further, a single dose of doxycycline treatment (4 μL of 10% w/w cream, i.e., 0.4 mg/incision) increased survival of *mntC*::Tn-infected larvae to 45% compared to only 7.5% survival in wild-type-infected ones (Fig. 1G). Together, knocking out *mntC* increased the sensitivity of USA300 to doxycycline *in vivo,* consistent with the *in vitro* SWF phenotype, demonstrating that MntC is a potential target for doxycycline adjuvant therapy for MRSA wound infections.

### A chemogenomic screen to uncover the mechanism of MRSA susceptibility to β-lactams in SWF

To investigate the increased sensitivity of USA300 to cefuroxime in SWF, we screened the NTML in SWF at 16 µg/mL of cefuroxime (4X the wild-type MIC in SWF) to identify susceptibility determinants. However, none of the mutants with greater than 25% relative growth at this supra-inhibitory concentration in SWF showed increased cefuroxime MIC in follow-up MIC assays (Fig. S7A-C). We also screened the NTML in MHB at 128 µg/mL of cefuroxime (1/8^th^ the wild-type MIC in MHB) to identify putative determinants of cefuroxime resistance, revealing 101 mutants with <8% growth at this sub-inhibitory concentration relative to their untreated controls (Fig. 2A-B). We hypothesized that mutants of determinants underlying cefuroxime sensitivity in SWF would be susceptible to cefuroxime in MHB, with no further sensitization in SWF relative to the parent strain. To that end, we tested the 101 mutants in dose-response and MIC assays. Nine mutants showed ∼128 to 256-fold sensitization to cefuroxime relative to the parent strain in MHB (i.e., showing MICs similar to or within 2-fold of that of USA300 in SWF) with relatively minimal MIC reduction relative to USA300 (≤4-fold) in SWF, revealing putative primary determinants of this phenotype (Fig. 2C). An additional 12 mutants showed a relatively more modest shift in cefuroxime MIC in MHB, potentially identifying secondary determinants of the phenotype (Fig. S8).

**Fig. 2.**
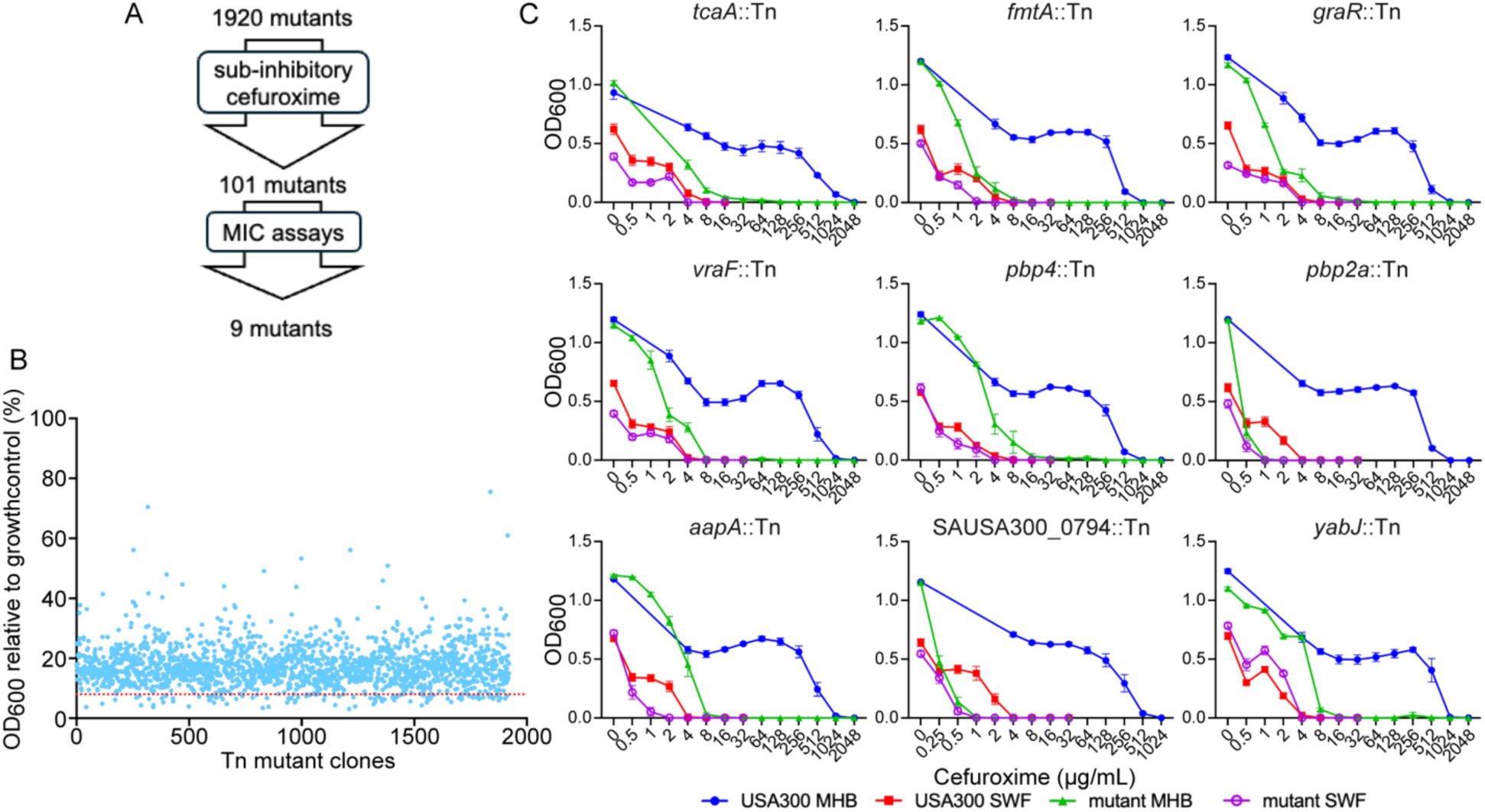
A chemogenomic screen identifies putative determinants of USA300 sensitivity to cefuroxime in SWF. A) The workflow of the chemogenomic screen and follow-up assays that led to the identification of putative determinants of cefuroxime sensitivity. B) A chemogenomic screen of the Nebraska Transposon Mutant Library with cefuroxime at 1/8^th^ MIC of the wild-type strain in MHB. C) MIC assays of putative determinants uncovered in the screen; n ≥ 6 from at least three independent experiments shown as mean ± SEM.

Six of the primary determinants (Fig. 2C) are localized to or involved in cell envelope homeostasis; their gene products are related to peptidoglycan synthesis [*pbp2a* and *pbp4*, encoding penicillin binding proteins (PBPs)], wall teichoic acid [WTA; *tcaA* and *fmtA*], and cell envelope stress response [*graR* and *vraF,* encoding members of the GraXSR-VraFG five-component response system] ^43–47^. Interestingly, some of the secondary determinants (Fig. S8) are involved in similar processes [MurA2, predicted to be involved in biosynthesis and degradation of peptidoglycan; RsbU and RsbV, involved in the regulatory cascade of the master transcriptional regulator σ^B^ that provides protection from stress and influences cell envelope composition ^48^] or localize at the membrane [SAUSA300_2297, SAUSA300_0073, SAUSA300_1785, SAUSA300_1484, SAUSA300_2315, and SpsA; predictions from ^49^]. The genes encoding the remaining three primary determinants are SAUSA300_0794 (encodes a TOPRIM domain-containing protein previously shown to be hypersensitive to the β-lactam amoxicillin ^50^), *aapA* (encodes a D-serine/D-alanine/glycine transporter), and *yabJ* (encodes a putative endo-RNase L-PSP). Importantly, most of the primary cell envelope-related determinants (Fig. 2C) are genetically interconnected as shown in synthetic lethal interaction analyses ^51^. Given that most mutants identified above harbour disruptions related to the cell envelope, which also comprises the β-lactam target (peptidoglycan), we then set out to characterize the primary cell envelope-related determinants and assess their implications for the unexpected β-lactam hypersensitivity of the MRSA USA300 in the wound-like medium.

### Decreased abundance of penicillin-binding proteins (PBPs) contributes to cefuroxime susceptibility in SWF

First, we investigated the potential role of alterations in PBPs in the SWF-specific cefuroxime susceptibility phenotype, especially given that PBPs are the main targets of β-lactams and that the primary resistance mechanism of MRSA is the low-affinity alternative PBP2a encoded by *pbp2a* or *mecA* ^52^. Based on *pbp2a*::Tn and *pbp4*::Tn cefuroxime susceptibility phenotypes (Fig. 2C), we hypothesized that USA300 cultured in SWF would have decreased abundance of these PBPs. To directly assess the abundance of PBPs in USA300, we isolated membrane fractions from cultures grown in MHB and SWF and then used a fluorescent penicillin (BOCILLIN^TM^ FL) to bind and detect PBPs resolved on SDS-PAGE. Quantification of band density identified a ∼25, ∼21, and ∼55 percent reduction in the relative abundance of PBP1, 2, and 4, respectively, in SWF compared to MHB (Fig. 3A, Fig. S9A, B). Notably, PBP1 and PBP2 are essential genes and therefore are not in the NTML ^46^. Also, BOCILLIN^TM^ FL labels all PBPs except PBP2a due to its decreased affinity for penicillin. Therefore, we assessed PBP2a abundance by Western blot using an anti-PBP2a-specific antibody ^53^, revealing a ∼20 percent decrease in the relative abundance of PBP2a in USA300 cultured in SWF relative to MHB (Fig. 3B, Fig. S9C, D). Together, the decreased abundance of the four PBPs (1, 2, 2a, and 4) in SWF is likely a major contributor to the decreased β-lactam susceptibility of USA300 under the wound-like condition.

**Fig. 3.**
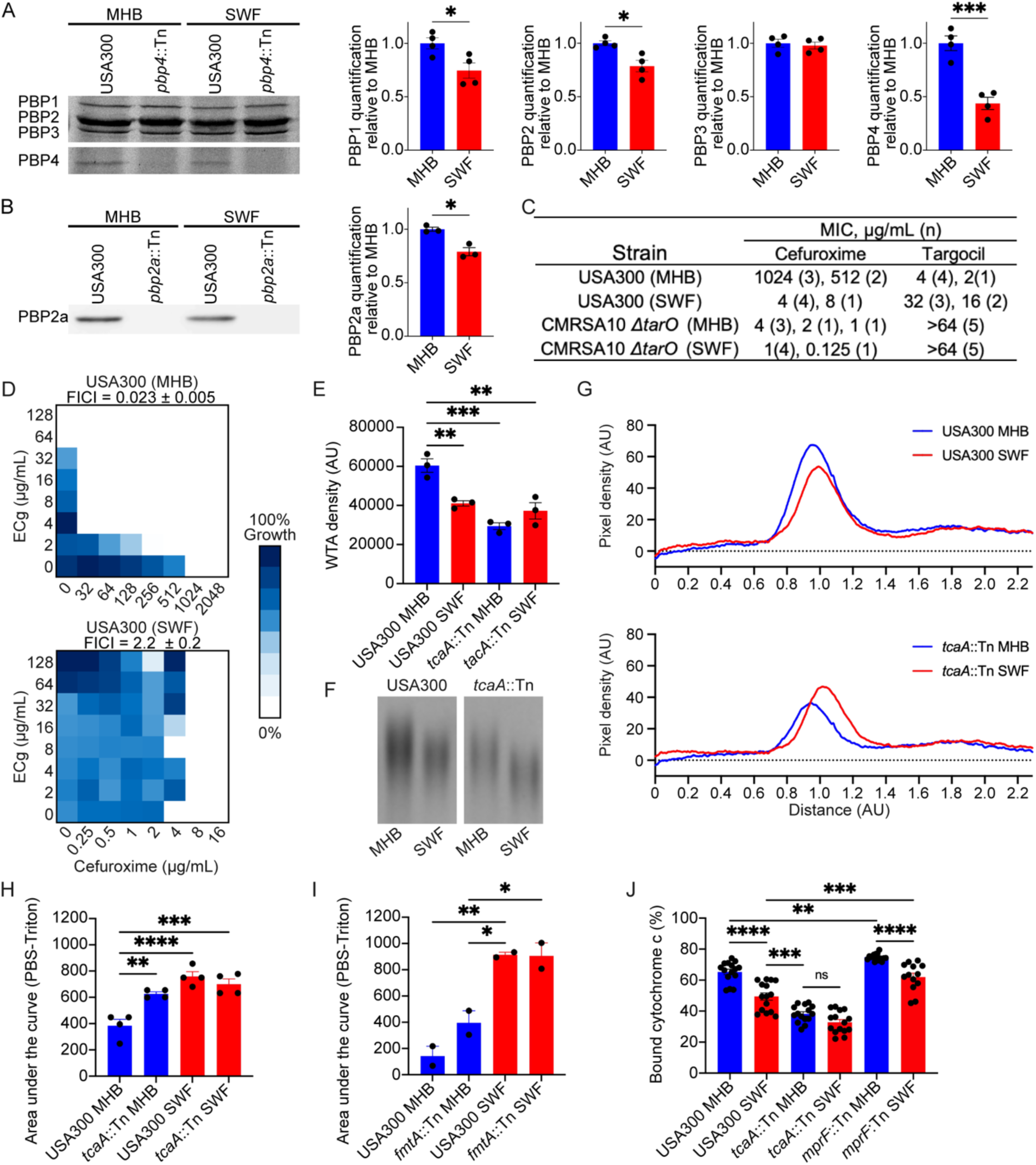
Decrease in the abundance of PBPs and wall teichoic acid in SWF contributes to USA300 cefuroxime sensitivity. A) Quantification of PBP1, 2, 3, 4 from USA300 cultured in SWF (red) relative to MHB (blue), n = 4 from two independent membrane isolations, shown as mean ± SEM and one representative BOCILLIN FL-labeled gel. All uncropped gels are shown in Fig. S9B) PBP2a quantification from Western blots, using an anti-PBP2a antibody, of membranes isolated from USA300 cultured in SWF (red) relative to MHB (blue), n = 3 from three independent membrane isolations, shown as mean ± SEM and one representative blot. All uncropped blots are shown in Fig. S9C) MIC of cefuroxime and targocil against USA300 or CMRSA10*ΔtarO* cultured in MHB or SWF, n = 5 biological replicates each performed in triplicate. D) Representative checkerboard assays of *S. aureus* USA300 of cefuroxime in combination with epicatechin gallate (ECg). n = 5. FICI shown as mean ± SEM. E) Relative quantity of PAGE-resolved crude WTA from *S. aureus* USA300 or *tcaA*::Tn cultured in MHB or SWF. n = 3 shown as mean ± SEM. F) Representative PAGE gel of WTA isolated from USA300 or *tcaA*::Tn cultured in MHB or SWF. Uncropped gel is shown in Fig. S10G) Relative band migration of WTA on PAGE represented as mean pixel density. n = 3 from three independent WTA isolations. H-I) Area under the curve (AUC) of lysis rates represented as PBS AUC – 0.02% Triton AUC comparing lysis rates of USA300 and *tcaA*::Tn (H; n=4) and *fmtA*::Tn (I; n=2) cultured from MHB or SWF. Shown as mean ± SEM. Full lysis curves and AUC calculation are shown in Fig. S10J) Percent cytochrome c binding to USA300 or mutants cultured in MHB or SWF, which infers relative cell charge. n = 15 or n = 13 for *mprF*::Tn SWF from 5 independent experiments, shown as mean ± SEM. Significant differences were determined by Welch’s t-test (A and B) and one-way ANOVA and the Tukey post hoc test (E, H, I, and J). p <0.0001 (****), p <0.001 (***), p <0.01 (**), and p <0.05 (*).

### Wall teichoic acid alterations contribute to USA300 sensitivity to cefuroxime in SWF

Next, we sought to evaluate changes in WTA, which plays a role in the coordination of peptidoglycan synthesis and β-lactam resistance ^54,55^. First, we sought to test *tarO,* which encodes a *N*-acetylglucosamine-1-phosphate transferase, the first committed step in WTA production ^56^. A *tarO* knockout is not within the NTML, and thus was not tested in our screen. A CMRSA10Δ*tarO* was ≥256-fold more susceptible to cefuroxime in MHB but only ∼4-fold more susceptible to the β-lactam in SWF relative to the wildtype (Fig. 3C), mirroring the phenotype of the primary determinants identified above (Fig. 2C). We then used targocil, an inhibitor of the WTA flippase TarG ^57^, to determine if culturing in SWF disrupts TarO or WTA biosynthesis. While targocil is lethal, knocking out genes encoding early steps in the WTA synthesis pathway (*tarO* and *tarA*) or inhibiting their products, which completely block WTA biosynthesis, antagonizes targocil ^58^. We observed an ∼8-fold increase in targocil MIC against wildtype USA300 when tested in SWF relative to MHB, raising the MIC up to 32 μg/mL (Fig. 3C). In comparison, the targocil MIC against the Δ*tarO* mutant was >64 μg/mL in both media (Fig. 3C), which together suggests that WTA biosynthesis is at least partially disrupted in SWF.

TarO inhibition has been associated with the delocalization of PBP2, leading to increased β-lactam sensitivity ^59^. Epicatechin gallate (ECg) induces partial delocalization of PBP2 from the division septum, mimicking the effects of TarO inhibition ^59^. As expected, ECg synergized with cefuroxime against the wild-type USA300 in MHB (FICI = 0.023 ± 0.005; Fig. 3D). However, such synergy was lost when we tested the same combination in SWF (FICI = 2.2 ± 0.2; Fig. 3D), suggesting PBP2 was already delocalized, potentially due to WTA biosynthesis disruption, under the wound-like condition.

Next, we directly evaluated the abundance of WTA in SWF and MHB (Fig. 3E-G, Fig. S10A). We tested both the wild-type USA300 strain and the *tcaA*::Tn mutant identified from the screen (Fig. 2), as *tcaA* encodes a teicoplanin resistance-associated protein involved in WTA retention and robustness of *S. aureus* peptidoglycan crosslinking ^43^. USA300 cultured in SWF showed a significant decrease in WTA abundance relative to MHB (Fig. 3E, F). The WTA abundance of the wild-type grown in SWF was comparable to that observed in the *tcaA*::Tn mutant cultured in MHB or SWF (Fig. 3E; no significant difference across these three conditions). Notably, the WTA bands from the wild-type cultured in SWF migrated slightly further on the PAGE relative to those from an MHB culture (Fig. 3F, G), suggesting a shorter WTA polymer. However, the shorter polymer length was not associated with TcaA, as the same occurred in the *tcaA*::Tn mutant (Fig. 3F, G).

WTA, its influence on PBP localization, and gene products associated with that polymer, such as FmtA (a teichoic acid D-Ala esterase) and TcaA, are involved in the structural integrity of the cell envelope, which can alter the bacterial autolysis ^43,44,60^. Therefore, we assessed the 0.02% Triton X-100-induced lysis of USA300, relative to knockouts of *fmtA* and *tcaA,* in MHB and SWF. Wild-type USA300 cells harvested from SWF cultures showed significantly faster Triton-induced lysis rates relative to those from MHB cultures (Fig. 3H, I, Fig. S10B-D). There was no further increase in lysis rates in SWF due to disrupting *tcaA* and *fmtA,* where both mutants showed no statistical difference in lysis rate compared to the wild-type cultured in SWF (Fig. 3H, I, Fig. S10B-D), suggesting WTA-associated gene products may have already been disrupted in SWF. In all cases, these observations are consistent with the alterations in WTA and PBP localization observed in SWF.

Next, we indirectly assessed cell surface charge, which is partly influenced by WTA retention, by quantifying binding to cytochrome c, a highly positively charged protein. A decreased abundance of WTA, a polyribitol phosphate polymer that generally imparts a strong negative charge to the surface ^61^, should reduce the overall net negative charge of the cell surface. USA300 cultured in SWF showed a significant decrease in binding to cytochrome c compared with MHB (Fig. 3J). We observed a decrease in relative cytochrome c binding to *tcaA*::Tn compared to that of USA300, but no difference in cytochrome c binding between *tcaA*::Tn cultured in MHB and SWF (Fig. 3J). These results align with the respective relative WTA abundance (Fig. 3E). Of note, other factors can also influence surface charge, such as MprF, a phosphatidylglycerol lysyltransferase, which decreases the net negative charge of the cell surface through lysyl modification of the cytoplasmic membrane, conferring resistance to cationic antimicrobials ^62^. While the cytochrome c binding of *mprF::Tn* in SWF relative to MHB followed the same trend observed for the wild-type strain, the lack of such modification in *mprF*::Tn increased the binding to cytochrome c relative to wild-type USA300 in each of the tested media, as expected (Fig. 3J). Taken together, the results in this section demonstrate WTA-related alterations that may contribute to the increased β-lactam susceptibility of the MRSA USA300 in the wound-mimetic medium.

### A lack of antibiotic-mediated induction of the GraXSR VraFG system in SWF

GraXSR VraFG is a five-component system that regulates the bacterial response to cell envelope stress, such as exposure to cationic antimicrobial peptides, through processes that include the DltABCD-mediated D-alanylation of teichoic acids and the MprF-dependent lysylination of phosphatidylglycerol ^63^. Notably, disruptions of GraR sensitize MRSA to β-lactams ^58^. *mprF* is positively regulated by GraXSR, and the system is strongly induced by colistin, a cationic antimicrobial peptide ^58^. Hence, we assessed the colistin-mediated induction of the GraXSR VraFG system in SWF compared to tryptic soy broth (TSB), a standard culture medium, by monitoring luciferase reporter expression under the control of the *mprF* promoter. As previously reported ^58^, we observed a colistin-mediated *graR*-dependent induction of the *mprF* promoter in TSB, where there was a significant increase in luciferase signal in the presence of colistin in the wild-type (Fig. 4A) but not in the Δ*graR* mutant (Fig. S11A-C). Such colistin-mediated *mprF* promoter induction was not observed in SWF (Fig. 4A); instead, a significant *graR*-independent repression of the *mprF* promoter at 3 hours by colistin was observed in both the wild-type (Fig. 4A) and the Δ*graR* mutant (Fig. S11D-F). To further assess the role of the GraXSR VraFG system in antibiotic response in SWF, we used MAC-545496, a GraR inhibitor previously shown to synergize with β-lactams in MHB ^58^. As expected, we observed a *graR*-dependent synergy between MAC-545496 and cefuroxime against USA300 in MHB (FICI = 0.036 ± 0.014; Fig. 4B; not observed against *graR*::Tn, FICI = 1.04 ± 0.47; Fig. S12). Such synergy was lost in SWF (FICI = 2.0 ± 0.0; Fig. 4B). Together, the lack of *mprF* promoter induction by colistin and loss of MAC-545496-cefuroxime synergy in SWF illustrate the lack of stress-mediated induction of GraR and its regulon, and hence the disruption of the GraXSR VraFG cell envelope stress response system, in the simulated wound fluid. Notably, such alteration in cell envelope stress response (Fig. 4) along with changes in cell envelope structures (the decreased abundance of several penicillin-binding proteins and wall teichoic acid; Fig. 3) of the MRSA USA300 under wound-like conditions can collectively result in the β-lactam sensitivity of MRSA in SWF (See Discussion Section and Fig. 7B).

**Fig. 4.**
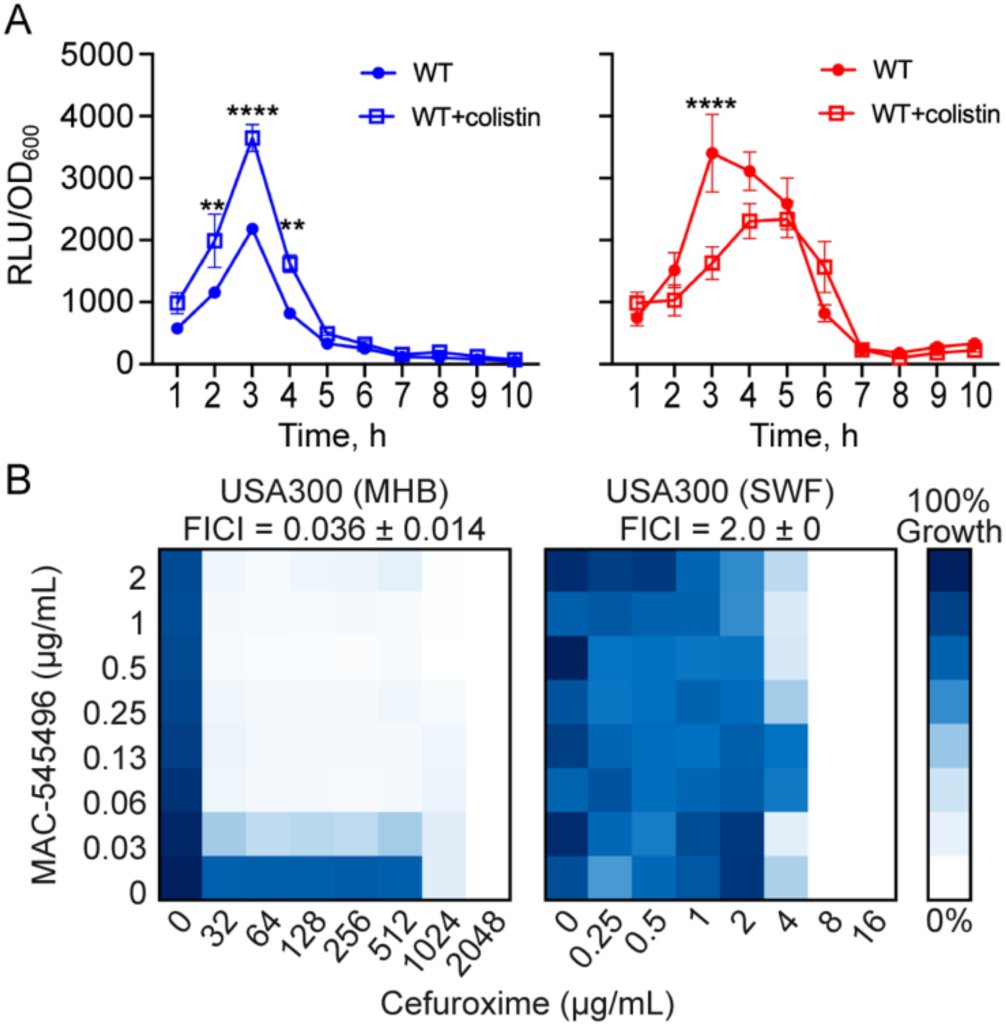
Disruption of GraXSR VraFG in SWF contributes to cefuroxime sensitivity. A) Monitoring the luciferase expression under the control of the *mprF* promoter in USA300 over 10 h in TSB (Blue) or SWF (Red) in the presence or absence of 1/32 MIC of colistin. n = 4 shown as mean ± SEM. Significant differences were determined between the with and without colistin conditions by two-way ANOVA and the Bonferroni post hoc test. p <0.0001 (****), p <0.001 (***), p <0.01 (**), p <0.05 (*). B) Representative cefuroxime and MAC-545496 checkerboard assays against *S. aureus* USA300. Dark blue represents high cell density, measured by OD_600_, n = 3, FICI shown as mean ± SEM.

### Ultrastructural, biochemical, and mechanical changes to USA300 grown in SWF

Next, we further assessed the cell and its envelope using atomic force microscopy (AFM) to compare cell size, surface topography, roughness, elasticity, and adhesion. We observed a significant increase in cell size (height and diameter) and surface roughness of USA300 cultured in SWF relative to that cultured in MHB (Fig. 5A-H, Fig. S13, 14). The surface of USA300 cells harvested from SWF culture was more compliant relative to those from MHB, requiring less force to deform (Fig. 5I). Adhesion values between the negatively charged silicon nitride AFM tip and the cell surface were greater for USA300 cells cultured in SWF relative to MHB (Fig. 5J), consistent with the cytochrome c binding assays showing that USA300 cultured in SWF are less negative than those in MHB (Fig. 3J). Together, these results provide further evidence for cell envelope-related alterations in simulated wound fluid, which underpins the increased β-lactam susceptibility.

**Fig. 5.**
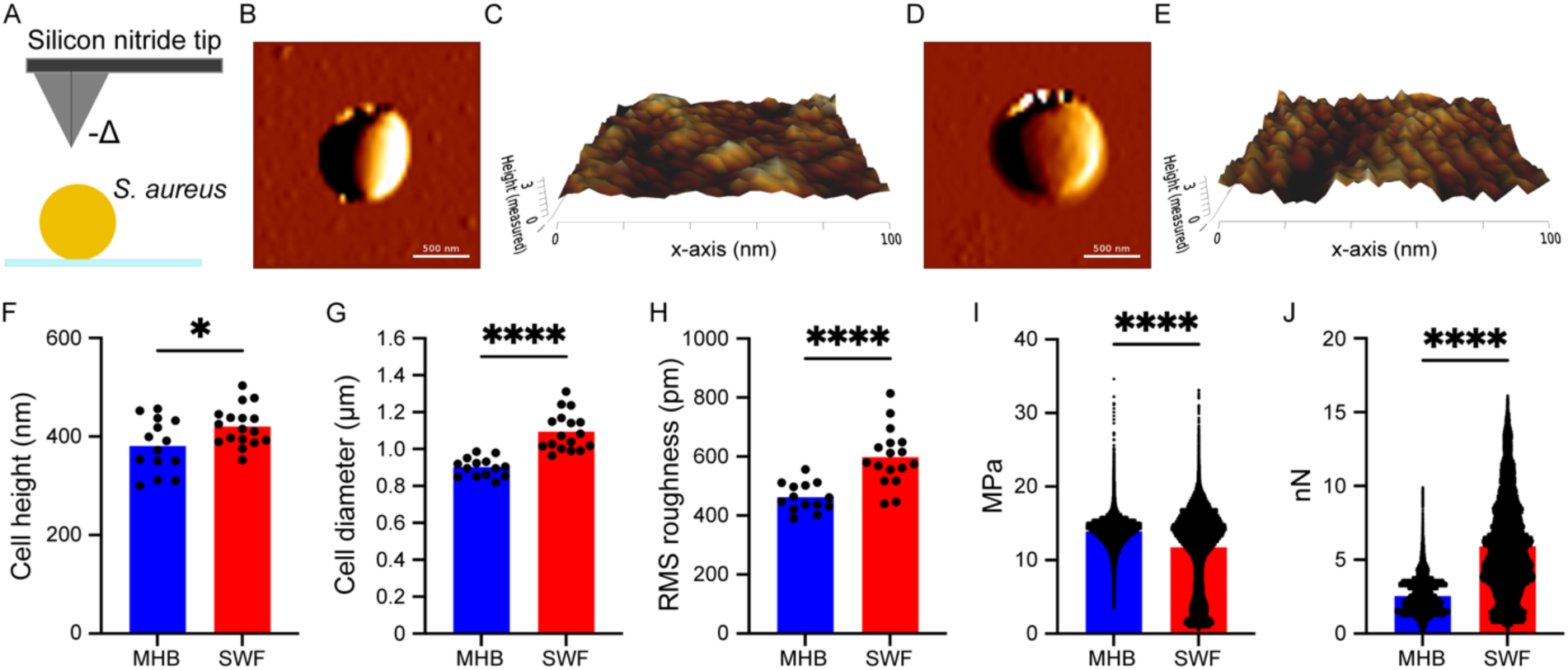
Atomic force microscopy of USA300 cultured in MHB and SWF. A) A schematic diagram of the AFM probe, B-E) Representative images (B, D) and 3D images of cell surface topography (C, E) of USA300 cultured in MHB (B, C) and SWF (D, E). All AFM images used for quantitative analyses are shown in Fig. S13 and S14F) Cell height of USA300 cultured in MHB and SWF. G) Cell diameter of USA300 cultured in MHB and SWF. H) Root mean squared (RMS) roughness of the flattest 100 nm x 100 nm region from a 0.4 x 0.4 µm scan area. I-J) Viscoelasticity and adhesion of USA300 cultured in MHB n = 91079 and SWF n = 118114. Significant differences between USA300 cultured in MHB and SWF were determined by Welch’s t-test. p <0.0001 (****), p <0.001 (***), p <0.01 (**), p <0.05 (*).

### Cefuroxime rescues USA300-infected Galleria larvae

Next, we sought to determine the efficacy of cefuroxime against USA300 in an *in vivo G. mellonella* incision wound infection model. Infection with wild-type USA300 treated with the vehicle control (Glaxal^TM^ base non-medicated cream), led to only 10% and 2.5% larval survival by days 3 and 6 post-infection, respectively (Fig.6). In contrast, a single dose cefuroxime treatment (4 μL of 40% w/w cream, i.e., 1.6 mg/incision) significantly increased the survival of USA300-infected larvae to ∼75% and 45% at the same timepoints post-infection (p<0.0001; Fig. 6). Together, the efficacy of cefuroxime against USA300 *in vivo* aligns with the *in vitro* susceptibility to that β-lactam antibiotic in SWF, demonstrating that this wound exudate-mimetic medium may be superior to MHB for predicting clinical outcomes of MRSA wound infections.

**Fig. 6.**
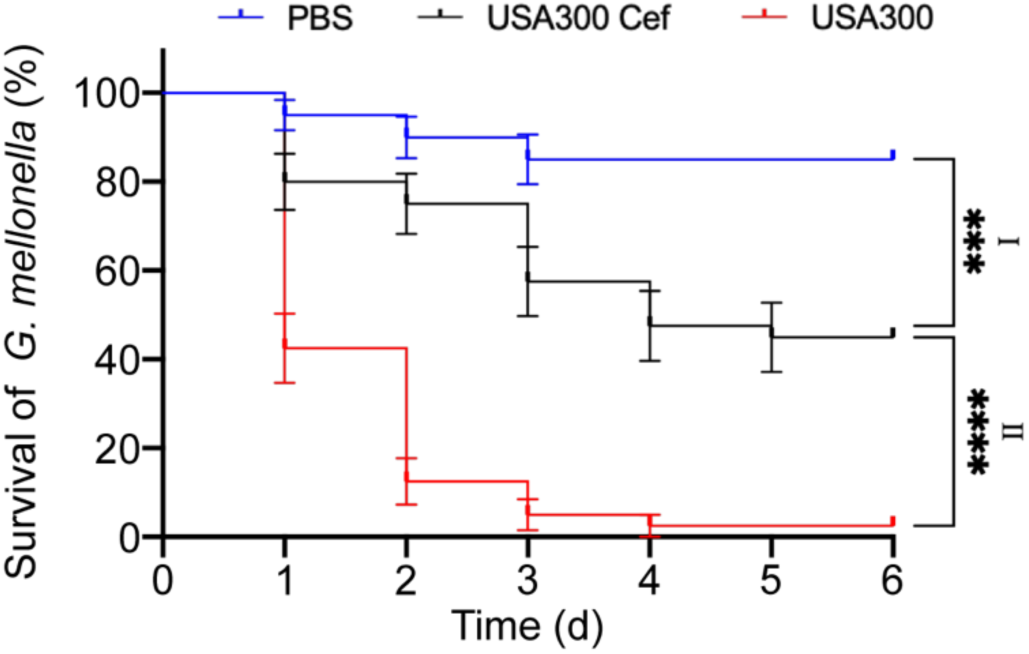
Cefuroxime increases larval survival *in vivo Galleria* wound infection. Kaplan-Meier survival curve of larvae shown as mean ± SE. PBS with vehicle treatment (blue), infection with USA300 and vehicle treatment (red), and infection with USA300 and cefuroxime treatment (black), n = 40 from 4 independent experiments. Significant differences were determined between (I; PBS and USA300 Cef) and (II; USA300 and USA300 Cef) by the Logrank (Mantel-Cox) test. p <0.0001 (****), p <0.001 (***), p <0.01 (**), p <0.05 (*).

## Discussion

Standard MHB-based AST assays have been widely used worldwide to guide antimicrobial therapy and study antimicrobial resistance. However, MHB does not accurately reflect the conditions at the site of infection ^64^, and the clinical outcome often does not correlate with standard AST results ^19–21^. Further, antibiotic discovery has traditionally relied on standard AST conditions, partly because of regulatory requirements to demonstrate efficacy under such conditions for approval ^65^, which has likely limited the suite of potential antimicrobial targets and bioactives. Therefore, there are emerging efforts to study bacterial responses and screen for new antimicrobials under infection-relevant conditions to bridge the gap between *in vitro* testing and *in vivo* outcomes [E.g., ^43,66–70^]. Herein, we studied the antibiotic responses of MRSA under wound-infection-mimetic conditions, revealing ∼67% discordance in AST results between simulated wound fluid and MHB. Similar discordances were observed in analyses using serum-containing and other eukaryotic cell culture media (e.g., RPMI) ^71,72^. We then sought to elucidate the mechanisms underlying the two most significant antibiotic susceptibility alterations in the MRSA USA300 strain in terms of fold-shift and potential clinical relevance, the unexpected resistance to tetracyclines and sensitization to β-lactams in SWF.

We revealed that the unexpected increase in resistance to first-, second-, and third-generation tetracyclines under wound-like conditions is mediated by MntC (Fig. 7A). MntC has not been previously known to confer tetracycline resistance, a role overlooked in standard culture conditions but detected in SWF. We observed an upregulation of *mntC* in SWF relative to MHB, independent of tetracycline exposure, suggesting MntC expression is induced by the culture condition rather than the antibiotic. Interestingly, the observed unexpected resistance to tetracyclines appears to corroborate reports of therapeutic failures with tigecycline (a tetracycline derivative) in soft-tissue, bloodstream, and other infections despite predicted efficacy ^73,74^. Also, the upregulation of *mntC*, which is highly conserved among *S. aureus*, in the serum-based SWF aligns with observations from *in vivo* studies and patient samples ^75–78^. MntC was only expressed *in vivo* or in the presence of serum, and rapidly upregulated in *S. aureus* murine infection ^75,76^. Anti-MntC antibodies protected against *S. aureus* in an infant rat model, and MntC could induce a protective immune response; hence, it was identified as a potential vaccine target ^75,76^. *mntC* (also referred to as *sitA*) had increased transcription in three healthy persistently colonized volunteers relative to *in vitro* conditions ^77^ and ranked among the first quartile of upregulated genes in pediatric skin and soft tissue infection (SSTI) and murine SSTI abscesses ^78^. As such, unexpected clinical failures of tetracyclines against MRSA may occur due to MntC overexpression *in vivo*. These observations, along with the increased survival of *mntC*::Tn-wound infected Galleria larvae, relative to wild-type, upon treatment with doxycycline (Fig. 1G), support that MntC could serve as an *in vivo*, clinically relevant drug target for new antibiotic adjuvants for tetracyclines. Interestingly, infection with an *mntC* knockout increased animal survival in a murine sepsis model, which was attributed to an increased susceptibility of the mutant to oxidative stress ^79^, aligning with the increased larval survival at day 1 post-infection with *mntC*::Tn relative to USA300 without treatment (Fig. 1G). These observations suggest that MntC may also serve as a target for stand-alone anti-virulence therapy.

**Fig. 7.**
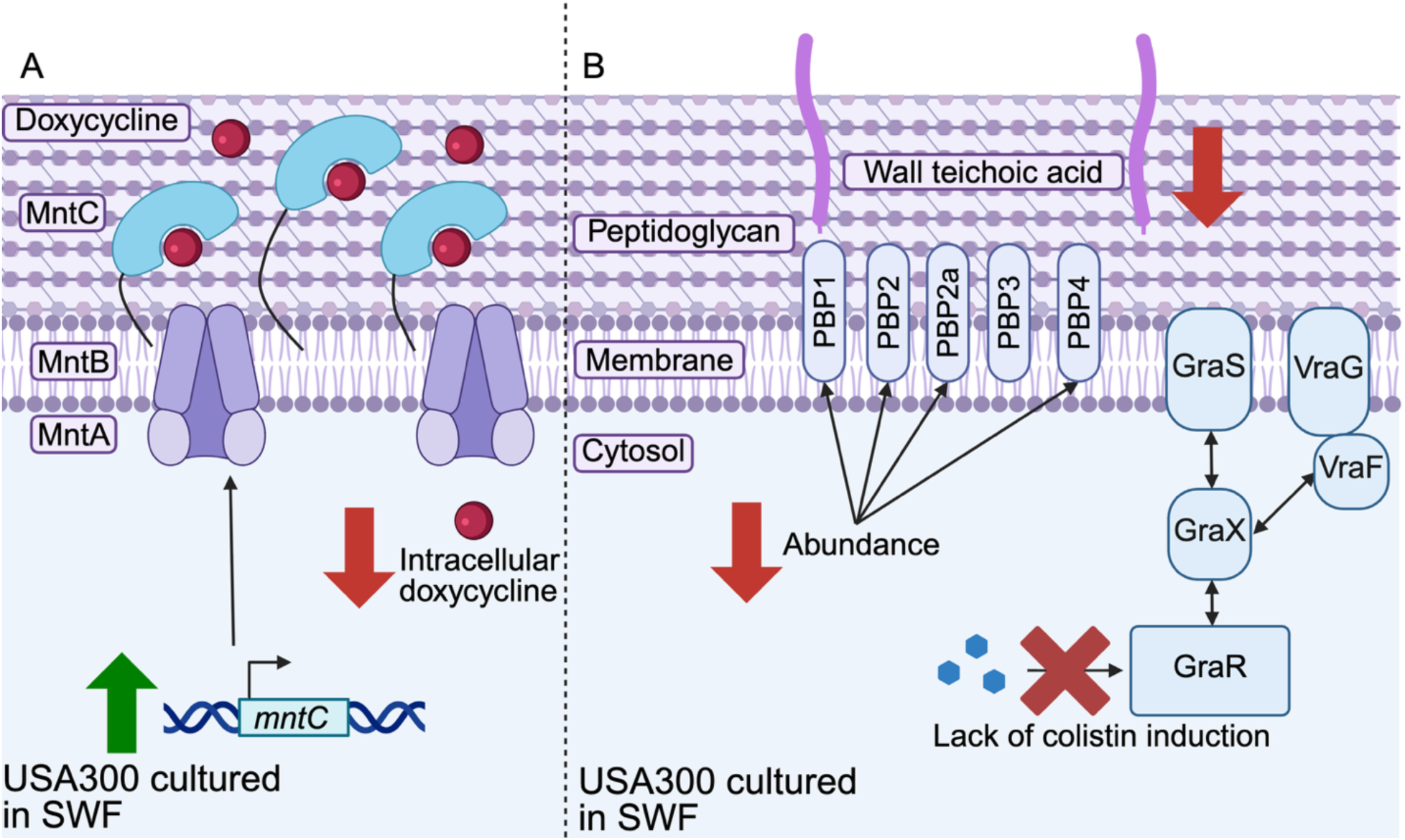
Summary of mechanisms underlying the altered susceptibility of MRSA USA300 to tetracyclines and β-lactams under wound infection-like conditions. A) Summary of alterations in USA300 that led to increased resistance to tetracyclines when cultured in SWF, depicting an increase in expression of MntC, which binds to doxycycline (a tetracycline), reducing its intracellular concentration. B) Summary of cell envelope-related remodelling in USA300 that increased susceptibility to β-lactams when cultured in SWF, displaying decreased abundance of penicillin binding proteins PBP1, 2, 2a, 4 and wall teichoic acid, along with a lack of the antimicrobial peptide colistin-mediated GraXSR VraFG induction. Created using Biorender.

MRSA are characterized by their resistance to β-lactam antibiotics. As such, the significant sensitization of tested MRSA strains from widely circulating lineages to representative β-lactams in the simulated wound fluid is counterintuitive but suggests that β-lactams may be a viable therapy for MRSA wound infections. *In vivo* results from the Galleria incision wound infection model aligned with the *in vitro* data, whereby the β-lactam cefuroxime improved the survival of MRSA USA300-infected larvae (Fig. 6). Interestingly, an observational study of 2096 pediatric skin and soft tissue infection patients at a community-acquired MRSA-endemic region in Philadelphia reported a therapeutic success of empiric β-lactam monotherapy comparable or even superior to that of MRSA-active antibiotics ^80^, suggesting that β-lactams likely cleared at least some MRSA infections, a regimen that standard AST would have contraindicated. Further, other recent reports have shown that adding β-lactams to the empirical therapy regimen improved bacterial clearance in MRSA bacteremia ^81–84^. Mechanistically, we revealed that cell envelope remodelling of the MRSA USA300 under wound-like conditions, involving a decreased abundance of penicillin binding proteins PBP1, 2, 2a, 4 and wall teichoic acid, along with a lack of antibiotic-mediated GraXSR VraFG induction (Fig. 7B), underpins the sensitivity of MRSA to β-lactams in SWF. Notably, various cell envelope changes were previously reported for *S. aureus* cultured in serum-containing media, altering responses to cell wall-targeting and membrane-active antibiotics ^43,66^. Together, our results and previous observations suggest that MHB-based AST may erroneously exclude β-lactams from therapeutic regimens, and that the wound exudate-like medium may be better than MHB at predicting β-lactam antibiotic efficacy. β-lactams are broad-spectrum, well-tolerated antibiotics that offer oral options ^85^; hence, accurately predicting their potency and recommending their use when possible would be valuable in the clinic.

Overall, host-mimetic media, such as SWF, may better predict bacterial resistance and correlate more closely with positive clinical outcomes. Uncovering antimicrobial susceptibility profiles discordant between infection-relevant (e.g., SWF) and standard (e.g., MHB) culture media may enhance our understanding of antibiotic therapeutic responses during infection. By revealing the mechanisms underlying the discordant phenotypes, we identified MntC as a potential target for new antimicrobials that may reverse resistance to currently available tetracycline antibiotics or suppress virulence *in vivo*. The phenotypes uncovered herein provide an *in vitro* screening platform in SWF for therapeutics targeting MntC. Notably, we have recently uncovered several natural bioactives that target MRSA planktonic and biofilm cells in simulated wound fluid, but not in standard media^86^, providing proof of concept for adjuvant discovery in SWF. Our AST results in the wound-like medium could also guide and optimize the use of currently available antibiotics.

## Limitations of the study

The *in vitro* AST and mechanistic studies under wound infection-mimetic conditions were conducted in the FBS-containing SWF^24^, which, while more representative of the wound exudates than MHB, may still miss some host-relevant components, some of which may be patient-specific, and lack the dynamic changes in composition and concentration that occur throughout the phases of wound healing. Importantly, wound exudates may vary in appearance and composition based on the etiology and specific conditions of the wound^25,87^. Hence, future testing in other simulated wound fluids^88,89^ with relatively minor compositional differences, potentially reflecting variations across patient groups and healing phases, may be complementary to this work. Additionally, the specific cues in SWF, representing potential host factors at wound sites (such as extracellular matrix and coagulation cascade components, including fibronectin, plasminogen, and fibrinogen that also bind to MntC^90^, metal ion availability, or other factors), and bacterial signalling pathways linking wound exudate microenvironmental exposure to cellular responses, including MntC upregulation and the coordinated cell envelope remodelling, are yet to be characterized, requiring further elucidation. Further, neither the *mntC*::Tn mutant nor the *mntB*::Tn mutant, with both *mntB* and *mntC* disrupted (Fig. S3C), fully accounted for the doxycycline MIC fold shift between MHB and SWF (Fig. 1A); a component of SWF inhibiting tetracyclines may have contributed to the unaccounted-for ∼2-fold increase in doxycycline MIC. The most significant antibiotic susceptibility phenotypes and the role of MntC in resistance to tetracyclines were tested *in vivo* in *G. mellonella*. While Galleria larvae mount both cellular and humoral innate immune responses closely resembling those of vertebrates ^41,42^ and their wound model recapitulates the hallmarks of trauma and infection observed in mammalian models ^32,33^, their immune system and surface tissues still differ from those of mammals. Thus, future studies should validate these Galleria results in *ex vivo* human skin and *in vivo* mammalian wound infection models.

## Resource availability

### Lead contact

- Further inquiries and requests should be directed to the lead contact, Omar El-Halfawy.

## Materials availability

- All reagents will be available upon request to the lead contact.

## Data and code availability

- All data reported in this study are available in the Main and Supplemental Tables and Figures included in this publication.
- This paper does not report original code.
- Any additional information required to reanalyze the data reported in this paper is available from the lead contact upon request.

## Acknowledgements

This work was funded by a New Frontiers in Research Funds exploration grant (NFRFE-2021-00398), a Saskatchewan Health Research Foundation establishment grant (6115), a Canadian Institutes of Health Research grant (202403PPE-522800), Canada Foundation for Innovation and Innovation Saskatchewan grants, and start-up funds from the Faculty of Science at the University of Regina to O.M.E. O.M.E. holds a Canada Research Chair in Chemogenomics and Antimicrobial Research (CRC-2024-00304). C.D.R. was supported by a Canadian Institutes of Health Research Canada Graduate Scholarship – Master’s (CGS-M) award. Work in the T.E.S.D. laboratory was supported by Natural Science and Engineering Research Council (NSERC DG 2024-06684) and Canada Foundation for Innovation grants to T.E.S.D. The Faculty of Graduate Studies and Research at the University of Regina provided partial support to A.M.

## Author contributions

Conceptualization: C.D.R., and O.M.E.; Formal analysis: C.D.R., and O.M.E.; Funding acquisition: O.M.E.; Investigation: C.D.R. and A.M.; Methodology: C.D.R., A.M., T.E.S.D., and O.M.E.; Project administration: O.M.E.; Resources: T.E.S.D. and O.M.E.; Supervision: T.E.S.D. and O.M.E.; Writing – original draft: C.D.R., and O.M.E.; Writing – review & editing: C.D.R., A.M., T.E.S.D., and O.M.E.

## Supplemental information

Document S1. Tables S1 and S2, Figures S1-S14, and supplemental references.

## STAR methods

### Key resources table

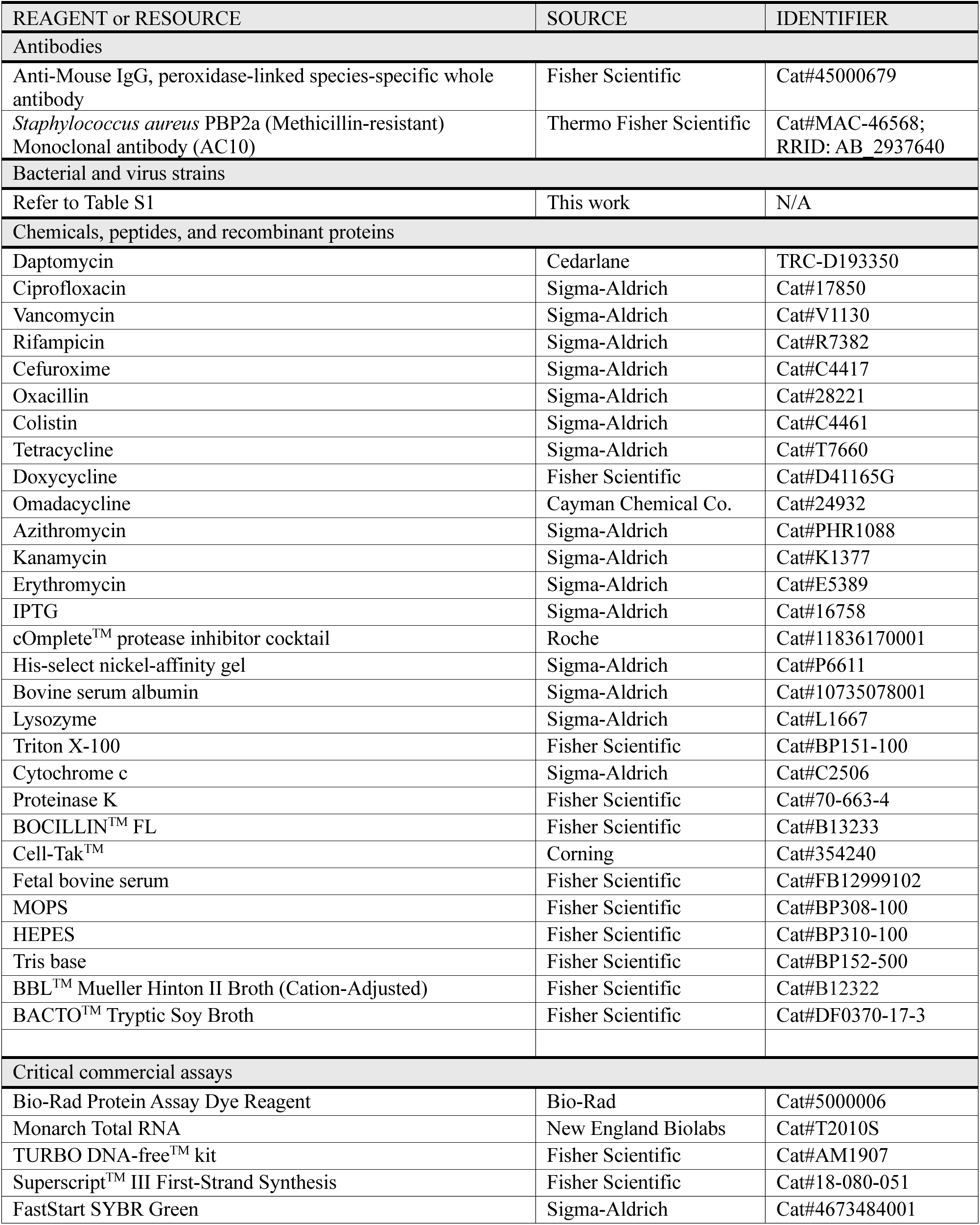
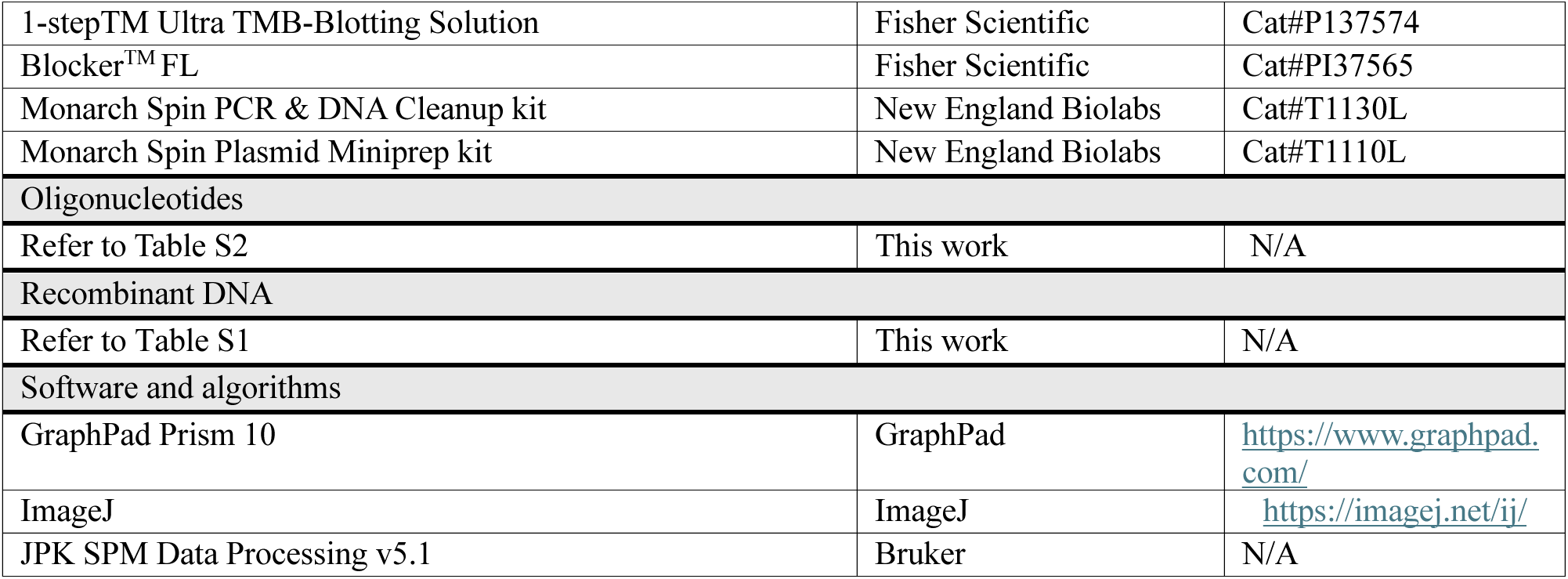

**STAR methods:** **Key resources table**
| REAGENT or RESOURCE | SOURCE | IDENTIFIER |
| --- | --- | --- |
| **Antibodies** |  |  |
| Anti-Mouse IgG, peroxidase-linked species-specific whole antibody | Fisher Scientific | Cat#45000679 |
| *Staphylococcus aureus* PBP2a (Methicillin-resistant) Monoclonal antibody (AC10) | Thermo Fisher Scientific | Cat#MAC-46568;<br>RRID: AB_2937640 |
| **Bacterial and virus strains** |  |  |
| Refer to Table S1 | This work | N/A |
| **Chemicals, peptides, and recombinant proteins** |  |  |
| Daptomycin | Cedarlane | TRC-D193350 |
| Ciprofloxacin | Sigma-Aldrich | Cat#17850 |
| Vancomycin | Sigma-Aldrich | Cat#V1130 |
| Rifampicin | Sigma-Aldrich | Cat#R7382 |
| Cefuroxime | Sigma-Aldrich | Cat#C4417 |
| Oxacillin | Sigma-Aldrich | Cat#28221 |
| Colistin | Sigma-Aldrich | Cat#C4461 |
| Tetracycline | Sigma-Aldrich | Cat#T7660 |
| Doxycycline | Fisher Scientific | Cat#D41165G |
| Omadacycline | Cayman Chemical Co. | Cat#24932 |
| Azithromycin | Sigma-Aldrich | Cat#PHR1088 |
| Kanamycin | Sigma-Aldrich | Cat#K1377 |
| Erythromycin | Sigma-Aldrich | Cat#E5389 |
| IPTG | Sigma-Aldrich | Cat#16758 |
| cOmplete™ protease inhibitor cocktail | Roche | Cat#11836170001 |
| His-select nickel-affinity gel | Sigma-Aldrich | Cat#P6611 |
| Bovine serum albumin | Sigma-Aldrich | Cat#10735078001 |
| Lysozyme | Sigma-Aldrich | Cat#L1667 |
| Triton X-100 | Fisher Scientific | Cat#BP151-100 |
| Cytochrome c | Sigma-Aldrich | Cat#C2506 |
| Proteinase K | Fisher Scientific | Cat#70-663-4 |
| BOCILLIN™ FL | Fisher Scientific | Cat#B13233 |
| Cell-Tak™ | Corning | Cat#354240 |
| Fetal bovine serum | Fisher Scientific | Cat#FB12999102 |
| MOPS | Fisher Scientific | Cat#BP308-100 |
| HEPES | Fisher Scientific | Cat#BP310-100 |
| Tris base | Fisher Scientific | Cat#BP152-500 |
| BBL™ Mueller Hinton II Broth (Cation-Adjusted) | Fisher Scientific | Cat#B12322 |
| BACTO™ Tryptic Soy Broth | Fisher Scientific | Cat#DF0370-17-3 |
| **Critical commercial assays** |  |  |
| Bio-Rad Protein Assay Dye Reagent | Bio-Rad | Cat#5000006 |
| Monarch Total RNA | New England Biolabs | Cat#T2010S |
| TURBO DNA-free™ kit | Fisher Scientific | Cat#AM1907 |
| Superscript™ III First-Strand Synthesis | Fisher Scientific | Cat#18-080-051 |
| FastStart SYBR Green | Sigma-Aldrich | Cat#4673484001 |
| 1-step<sup>TM</sup> Ultra TMB-Blotting Solution | Fisher Scientific | Cat#P137574 |
| Blocker<sup>TM</sup> FL | Fisher Scientific | Cat#PI37565 |
| Monarch Spin PCR & DNA Cleanup kit | New England Biolabs | Cat#T1130L |
| Monarch Spin Plasmid Miniprep kit | New England Biolabs | Cat#T1110L |
| Oligonucleotides |  |  |
| Refer to Table S2 | This work | N/A |
| Recombinant DNA |  |  |
| Refer to Table S1 | This work | N/A |
| Software and algorithms |  |  |
| GraphPad Prism 10 | GraphPad | https://www.graphpad.com/ |
| ImageJ | ImageJ | https://imagej.net/ij/ |
| JPK SPM Data Processing v5.1 | Bruker | N/A |

### Strains and reagents

Table S1 lists the strains and plasmids, and Table S2 lists the primers used in this study. Bacteria were grown in cation-adjusted Mueller-Hinton broth (MHB) or simulated wound fluid (50% Fetal bovine serum from Fisher Scientific, 50% Maximum recovery diluent (MRD; 1g/L peptone, 8.5g/L NaCl) at 37 °C, unless otherwise specified.

### Antimicrobial Susceptibility Testing (AST)

MICs were determined by broth microdilution according to CLSI guidelines ^93^. The assays were conducted in microtiter plates with a final inoculum adjusted to an OD_600_ of 0.001, prepared by the colony resuspension method. The plates were then incubated at 37°C with shaking, and bacterial growth was determined by measuring the OD_600_ after 24 h. When MIC values were not reached within the tested concentration range, the highest tested concentration was considered half the MIC. The fractional inhibitory concentration indices (FICI) were calculated as FICI = (MIC_A_ in combination/MIC_A_ alone) + (MIC_B_ in combination/MIC_B_ alone). FICI values ≤0.5 were interpreted as synergy, >0.5 to ≤4.0 as additive or indifferent, and >4.0 as antagonistic.

### Nebraska transposon mutant library (NTML) screen

Overnight cultures of the NTML ^38^ were prepared in 384-well plates using Biomatrix BM6-BC (S&P Robotics Inc.) in MHB with 5 µg/mL erythromycin (Sigma-Aldrich). The following day, these overnight cultures were used to inoculate MHB or SWF with or without doxycycline (Fisher Scientific) or cefuroxime (Sigma-Aldrich), at the concentrations indicated in the results section, using the BM6-BC. All 384-well plates were incubated at 37 °C and 600 RPM, and the OD_600_ was read after 24 h.

### Dose-response assay

Selected NTML strains were re-arrayed and inoculated into 96-well plates in MHB containing 5 µg/mL erythromycin using the BM6-BC. The following day, 96-well plates with varying antibiotic concentrations in MHB and SWF were inoculated using the BM6-BC. All plates were incubated at 37 °C and 600 RPM, and the OD_600_ was read after 24 h.

### Genetic complementation

To complement *mntB*::Tn and *mntC*::Tn, PCR fragments were amplified using USA300 gDNA as template and primer pairs (listed in Table S2): mntB_compF3_AvrII and mntB_compR2_BamHI for *mntB*, mntB_compF3_AvrII and mntC_compR_BamHI for *mntBC*, and mntC_compF3_AvrII and mntC_compR_BamHI for *mntC*. Fragments and pKK22-Pfba were digested with AvrII and BamHI (New England Biolabs), and the plasmid was alkaline phosphatase-treated (using Antarctic phosphatase, New England Biolabs), followed by ligation with T4 DNA ligase (New England Biolabs) to generate the complementation vectors, pKK22-Pfba-*mntB*, pKK22-Pfba-*mntBC*, and pKK22-Pfba-*mntC*. Complementation and control vectors were transformed into electrocompetent *S. aureus* strains by electroporation (Bio-Rad) ^94^.

### Doxycycline accumulation assay

Cultures were started from a single colony and incubated overnight in MHB or SWF at 37 °C with shaking, then sub-cultured by diluting to an OD_600_ of 0.1 in fresh media and further incubated for 6 h. Cells were harvested from the sub-cultures and resuspended at an OD_600_ of 1 in MHB or SWF. Doxycycline was added to 1 mL of these OD-adjusted cultures at a final concentration of 5 µg/mL, and the cultures were incubated for 30 min at 37 °C. Cells were pelleted, resuspended in 1 mL of fresh culture media, and incubated for 15 min at 37 °C. The cells were then washed twice with PBS and resuspended in 0.5 mL of PBS, followed by dispensing of 100 µL aliquots into 96-well black plates in quintuplicate and read at λ_ex_ = 400 nm / λ_em_ = 530 nm. The bacterial suspensions were also transferred to a clear 96-well plate for OD_600_ measurement. All centrifugation steps were at 6,000 × g for 1 min.

### Protein overexpression and purification

*mntC* was cloned into pET28a(+) and transformed into CaCl_2_ chemically competent *E. coli* BL21 by heat shock. Bacterial cultures were incubated at 30 °C and 250 RPM until an OD_600_ of ∼0.8 was reached, then induced with 0.5 mM IPTG and further incubated for 5 h at 30 °C. Cells were harvested, resuspended in lysis buffer [50 mM sodium phosphate, 500 mM NaCl, cOmplete^TM^ protease inhibitor cocktail (Roche), pH 8.0], and lysed using Lysing Matrix B (MP Biomedicals) and the FastPrep-24^TM^. The supernatants were separated from the insoluble fractions by centrifugation at 10,000 × g for 15 min at 4 °C. His-tagged proteins were batch purified from the supernatants using His-select nickel-affinity gel (Sigma-Aldrich), washed three times with wash buffer (50 mM sodium phosphate, 500 mM NaCl, 5 mM imidazole, pH 8.0), and eluted with elution buffer (50 mM sodium phosphate, 500 mM NaCl, 250 mM imidazole, pH 8.0), followed by detection on a 12% acrylamide gel (Bio-Rad) stained with PageBlue^TM^ (Fisher Scientific). The eluted protein was dialyzed using 10 kDa SnakeSkin^TM^ dialysis tubing (Fisher Scientific) into 20 mM HEPES, pH 7.4, 100 mM NaCl buffer, then quantified using the Bradford method with protein assay dye reagent concentrate (Bio-Rad).

### Doxycycline binding assay

Purified MntC, bovine serum albumin (BSA), and lysozyme solutions in dialysis buffer (20 mM HEPES, pH 7.4, 100 mM NaCl) were each incubated at 37 °C with doxycycline in the dark for 4 h at a final volume of 200 µL and final concentration of 32 µM of protein and 16 µM of doxycycline in 96-well black plates. Wells containing only 200 µL of buffer with 16 µM of doxycycline and 200 µL of buffer with 32 µM of protein were used for background subtraction of fluorescent signal. Plates were then read at λ_ex_ = 390 nm / λ_em_ = 550 nm.

### RNA extraction

USA300 cultures were grown overnight and diluted to an OD_600_ of 0.05 in MHB or SWF. Cells were grown at 37 °C to an OD_600_ of 0.6, then treated with doxycycline (8 µg/mL) or vehicle control (ultrapure water) for 30 minutes. Aliquots (6 mL) of the cultures were collected, centrifuged at 3,320 × g for 10 min. RNA was extracted using Monarch Total RNA miniprep (New England Biolabs), followed by treatment with Invitrogen Turbo DNA-free DNase (Fisher Scientific) following the manufacturer’s protocol. RNA purity was assessed by measuring the ratio of absorbance at 260 nm to 280 nm (values obtained ∼2.0).

### cDNA synthesis and RT-PCR

RNA was converted to cDNA with Superscript™ III First-Strand Synthesis System (Fisher Scientific) following the manufacturer’s protocol. RT-PCR was performed using the QuantStudio^TM^ 5 Real-Time PCR (Thermo Fisher Scientific) with FastStart SYBR Green (Roche). Fold change in gene expression was calculated using the Pfaffl Method ^95^ relative to *arcC* and 16s rRNA.

### *Galleria mellonella* incision wound infection model

*G. mellonella* larvae weighing ∼300 mg were selected, anesthetized by chilling at 4 °C for 10 minutes, and kept on ice during incision and treatment application. Incisions spanning the third segment from the posterior end of the larvae were made parasagittally on the dorsal side. Larvae were inoculated with 2 µL of inoculum (0.0001 OD_600_/mg of larva weight). After 30 min, 4 µL of a 10% w/w doxycycline compounded in Glaxal^TM^ Base, 40% w/w cefuroxime compounded in Glaxal^TM^ Base, or vehicle control (water in Glaxal^TM^ Base) was applied to the inoculated wounds. Worms were then incubated at 37 °C and monitored for survival for 6 days. Of note, research ethics approval is not required for the use of this invertebrate model.

### Lysis Assay

Overnight cultures were diluted to an OD_600_ of 0.1 and grown to an OD_600_ of 0.8. Cells were washed and resuspended in PBS or PBS containing 0.02% Triton X-100 (Fisher Scientific) at a cell density of 1 OD/mL. The bacterial suspensions were dispensed into 200 µL aliquots in triplicate in 96-well plates, incubated at 37 °C with shaking in the plate reader, and their OD_600_ was read every 15 min for 6 h.

### Cytochrome c binding assay

The assay was performed as previously described ^96^. Briefly, overnight cultures were adjusted to an OD_600_ of 1, and 8 mL were pelleted, washed twice in MOPS buffer (20 mM, pH 7.0), and resuspended in a total volume of 200 µL MOPS. A 50 µL aliquot of a 2.5 mg/mL cytochrome c solution (Sigma-Aldrich) was added to the 200 µL resuspended culture and incubated at room temperature for 10 min. The samples were then pelleted at 16,000 × g for 1 min, and 200 µL of the supernatants were dispensed into 96-well plates, and their absorbance was read at 530 nm (A_530nm_). *mprF*::Tn was used as a positive control.

### Bioluminescence assay

The luciferase expression assay was conducted as previously described ^58^, except that bacteria were cultured in either TSB or SWF. Briefly, USA300 (pGYlux::*mprF*), USA300 (pGYlux::*ilvD*), Δ*graR* (pGYlux::*mprF*), and USA300 with control vector pGYlux were grown overnight in TSB and SWF with 10 µg/mL chloramphenicol. The overnight cultures were then diluted to an OD_600_ of 0.01 in 10 mL of TSB or SWF with and without 1/32^nd^ MIC of colistin and grown at 37 °C with shaking. Each hour, 100 µL was plated into a white 96-well plate in triplicate, and relative light units (RLU) were measured before transferring to a clear 96-well plate to measure OD_600_.

### WTA isolation

Crude WTA was isolated as previously described ^97^, with minor modifications to normalize the quantity of bacteria prior to WTA isolation. Briefly, overnight cultures in either MHB or SWF were washed once with buffer 1 (50 mM MES, pH 6.5) and volumes corresponding to an OD_600_ of 5 were centrifuged at 5,000 × g, resuspended in buffer 2 [4% (w/v) SDS, 50 mM MES, pH 6.5], and boiled for 1 h, followed by washing in buffer 1 and transferring into 1.5 mL microcentrifuge tubes. Pellets were then washed with 1.4 mL of buffer 2, then with 1.4 mL of buffer 3 (2% NaCl, 50 mM MES, pH 6.5), followed by 1.4 mL of buffer 1 with all spins at 16,000 × g for 1 min. Pellets were resuspended in 1 mL of digestion buffer [20 mM Tris-HCl, pH 8.0, 0.5% (w/v) SDS] with the addition of 10 µL of 2 mg/mL proteinase K. Samples were digested at 50 °C for 4 h with shaking (1000 rpm). Pellets were then washed with 1.4 mL of buffer 3, followed by three 1.4 mL washes of ultrapure water. The pellets were then resuspended in 1 mL of 0.1 M NaOH and incubated at 20 °C for 16 h with shaking at 1000 rpm in an infors HT Multitron incubator (3 mm throw diameter). The samples were then centrifuged at 16,000 × g for 1 min, and 750 µL of the supernatants were transferred to a new microcentrifuge tube. Subsequently, 250 µL of 1 M Tris-HCl, pH 7.8, was added to neutralize the reaction.

### WTA PAGE

Aliquots (15 µL) of crude WTA were mixed with 5 µL of 4x loading dye (50% glycerol with bromophenol blue), and 10 µL was loaded and separated on tricine polyacrylamide gels at 40 mA ^97^ until the dye front had reached the bottom of the gel. Gels were washed three times with ultrapure water, then stained with Alcian Blue 8GX (Fisher Scientific, 1 mg/mL) overnight, and destained with ultrapure water until the bands became visible.

### Membrane preparation

Cultures grown overnight at 37 °C with shaking were diluted to an OD_600_ of 0.05 and then grown to an OD_600_ of 0.8. The cultures were pelleted, resuspended in PBS, and lysed using Lysing Matrix B (MP Biomedicals) and the FastPrep-24^TM^. Cell debris was pelleted at 10,000 × g for 30 min, and the supernatant was centrifuged at 40,000 rpm for 1h to pellet the membrane, which was then resuspended in PBS, and its protein content was quantified by the Bradford method using protein assay dye reagent concentrate (Bio-Rad).

### BOCILLIN^TM^ FL assay

Membrane fractions (equivalent to 30 µg of protein content) from USA300 and *pbp4*::Tn cultured in MHB and SWF were incubated with 20 µM of BOCILLIN^TM^ FL (Fisher Scientific) at 37 °C for 10 min, followed by the addition of 4x Laemmli buffer and boiling for 5 min. Membrane samples were resolved on 10% SDS-PAGE running at 80V for 30 min, followed by 120V for 2.25 h. Gels were then visualized using the Typhoon FLA 7000 gel scanner (GE Healthcare), and the band intensities were subsequently quantified with ImageJ. PBP quantifications were background-subtracted by membrane-free BOCILLIN^TM^ FL lanes.

### PBP2a western blot

Membrane fractions (equivalent to 10 µg of protein content) from USA300 and *pbp2a*::Tn cultured in MHB and SWF were resolved on 10% SDS-PAGE, running at 80 V for 30 min, followed by 120 V for 2.25 h. Protein bands were transferred to nitrocellulose film using a Mini Trans-Blot® Cell (Bio-Rad) at 100 V for 2 h. The nitrocellulose membrane was then blocked overnight at 4 °C with Blocker^TM^ FL (Thermo Scientific). The primary antibody, mouse anti-*S. aureus*-PBP2a (Thermo Fisher Scientific) (1:1000 dilution in Blocker^TM^ FL), was incubated for 2 h at room temperature with rocking. The secondary antibody, a horseradish peroxidase-conjugated sheep anti-Mouse IgG (Thermo Scientific) (1:1000 dilution in Blocker^TM^ FL buffer), was incubated for 2 h at room temperature with rocking. The blot was developed with 1-Step^TM^ Ultra TMB-Blotting Solution (Thermo Scientific) for 10 min, then visualized on the Gel Doc^TM^ EZ imager (Bio-Rad), and band intensities were subsequently quantified with ImageJ.

### Atomic force microscopy (AFM) imaging

AFM (Nanowizard 4; JPK Instruments, Berlin, Germany) in quantitative imaging (QI™) mode ^98,99^ was used to analyze cell surface topography and mechanics of USA300 cells cultured in MHB and SWF. Overnight cultures of USA300 in MHB and SWF were diluted to an OD_600_ of 0.05 and grown to an OD_600_ of 0.6. Cells were then washed three times in PBS and resuspended to an OD_600_ of 0.125 and deposited onto clean, Cell-Tak^TM^ (Corning®)-coated coverslips ^100^, incubated for 30 min at room temperature followed by rinsing with ultrapure water three times, air dried (30 min) at room temperature, and imaged using non-conductive silicon nitride cantilevers (Multi-cantilever Tip (MLCT), D cantilever, nominal k = 0.03 N/m and tip radius 20 nm; Bruker, USA). Cantilever spring constant (0.028 N/m) was calibrated using the thermal noise method ^101^. QI™ force curves at each pixel of a 128 × 128 raster scan were collected using a Z-length of 1000 nm, setpoint of 2 nN, and a raster scan of 100 µm/s, along with a subset of force curves within a 0.4 × 0.4 nm square in the center of the cell. Force curves were batch-processed using JPK SPM Data Processing software (version 5.1, JPK, Berlin, Germany), and histogram data were exported to Excel.

### Statistical analysis

All statistical analyses were calculated using GraphPad Prism 10. The statistical test used for each assay is indicated in the figure captions.

