## Supplemental Figures and Tables for "Uncovering the mechanisms of clinically-relevant altered antibiotic responses of *Staphylococcus aureus* under wound infection-mimetic conditions"

**Running title:** Altered antibiotic responses of *S. aureus* under wound-like conditions

**Supplemental information: Document S1**

- **Tables S1 and S2**
- **Figures S1-S14**
- **Supplemental references**

**Table S1. Strains and plasmids used in this study**

| Bacterial Strain | Description | Reference |
|---|---|---|
| *Staphylococcus aureus* | | |
| USA300 | *S. aureus* USA300 LAC isolated in 2002 from a skin and soft tissue infection of an inmate in the Los Angeles County Jail in California, USA.; hypervirulent community-associated MRSA; cured of antibiotic resistance plasmid; also known as JE2; parent of the NTML. | Lab stock* |
| RN4220 | Restriction endonuclease deficient lab strain | [1] |
| CMRSA 1 | HA MRSA USA600 lineage strain, ST45, CC45, SCC*mec* type II | Lab stock [2] |
| CMRSA 2 | HA MRSA USA100/800/New York lineage strain, ST5, CC5, SCC*mec* type II | Lab stock [2] |
| CMRSA 3 | HA MRSA strain, ST241, CC8, SCC*mec* type III | Lab stock [2] |
| CMRSA 4 | HA MRSA USA200/EMRSA16 lineage strain, ST36, CC30, SCC*mec* type II | Lab stock [2] |
| CMRSA 5 | HA MRSA USAA500 lineage strain, ST8, CC8, SCC*mec* type IV | Lab stock [2] |
| CMRSA 6 | HA MRSA strain, ST239, CC8, SCC*mec* type III | Lab stock [2] |
| CMRSA 7 | CA MRSA USA400/MW2 lineage strain, ST1, CC1, SCC*mec* type IV | Lab stock [2] |
| CMRSA 8 | HA MRSA EMRSA15 lineage strain, ST22, CC22, SCC*mec* type IV | Lab stock [2] |
| CMRSA 9 | HA MRSA strain, ST8, CC8, SCC*mec* type II | Lab stock [2] |
| CMRSA 10 | CA MRSA USA300 lineage strain, ST8, CC8, SCC*mec* type IV | Lab stock [2] |
| USA300*ΔgraR* | USA300 strain with *graR* deletion | [3] |

| | | |
|---|---|---|
| NTML | Nebraska Transposon Mutant Library Screening Array: 1920 *S. aureus* subsp. *aureus* USA300 JE2, transposon (Tn) mutants arrayed in five 384-well microtiter plates. Erm$^R$ | [4*] |
| CMRSA10*ΔtarO* | CMRSA10 strain with *tarO* deletion | [5] |
| USA300*ΔsbnB* | USA300 with *sbnB* deletion | [6] |
| *mntB*::Tn pKK22-Pfba-*mntB* | CDR24: USA300 *mntB*::Tn with pCR47 | This study |
| *mntB*::Tn pKK22-Pfba-*mntBC* | CDR27: USA300 *mntB*::Tn with pCR48 | This study |
| *mntC*::Tn pKK22-Pfba-*mntC* | CDR26: USA300 *mntC*::Tn with pCR37 | This study |
| *Escherichia coli* | | |
| DH5αλpir | Lab strain engineered for transformation efficiency, containing the *pir* genes | Lab stock** |
| BL21 | Lab strain with T7 RNA polymerase for expressing recombinant proteins | Lab stock |

| Plasmid | Description | Reference |
|---|---|---|
| pGYlux::*ilvD* | pGYlux with *ilvD* promoter cloned | [3] |
| pGYlux::*mprF* | pGylux with *mprF* promoter cloned | [7] |
| pKK22-Pfba | pKK22 [8]** with *fba* promoter cloned | Lab stock |
| pKK22-Pfba-*mntB* | pCR47: Vector for complementation of *mntB*::Tn | This study |
| pKK22-Pfba-*mntBC* | pCR48: Vector for complementation of *mntB*::Tn | This study |
| pKK22-Pfba-*mntC* | pCR37: Vector for complementation of *mntC*::Tn | This study |
| pET28a(+) | IPTG-inducible expression vector with a 6X His tag | Lab stock |
| pET28a(+)-*mntC* | pCR20: pet28a(+) with *mntC* without the sequence encoding the entire N-terminal lipoprotein signal sequence (18 amino acids, including the lipobox Cys residue) as described in [9] | This study |

* Provided by the Network on Antimicrobial Resistance in *Staphylococcus aureus* (NARSA) for distribution by BEI Resources, NIAID, NIH.
** Contributed by Dr. J. L Bose for distribution by BEI Resources, NIAID, NIH.
HA: Hospital-acquired; CA: Community-acquired; CC: Clonal cluster; ST: Sequence type

**Table S2. Primers used in this study**

| Primer Name | Sequence |
|---|---|
| JE2_mntC_F_qPCR | GGTGGAGACAACGTCGATATTC |
| JE2_mntC_R_qPCR | CCAACCGTTACCAGTCTCTAAAT |
| JE2_16s_F_qPCR | GTGGAGGGTCATTGGAAACT |
| JE2_16s_R_qPCR | CACTGGTGTTCCTCCATATCTC |
| JE2_arcC_F_qPCR | CGGTAATGCGATACAGACAACA |
| JE2_arcC_R_qPCR | CGCGCTGGTGAATCAAATAAAG |
| mntCpet28aFTNdeI | TATACATATGATGGGTACTGGTGGTAAACAAAGCA |
| mntCpet28aRTBamHI | TATAGGATCCTTATTTCATGCTTCCGTGTACAGT |
| mntB_compF3_AvrII | TATACCTAGGATGTTAGAGTTTGTCGAACATTTATTTACATATC |
| mntB_compR2_BamHI | ATATGGATCCTCATGTTAAACTTCCTCGTTTCTTTCTATTCG |
| mntC_compF3_AvrII | TATACCTAGGATGAAAAAATTAGTACCTTTATTATTAGC |
| mntC_compR_BamHI | TATAGGATCCTTATTTCATGCTTCCG |

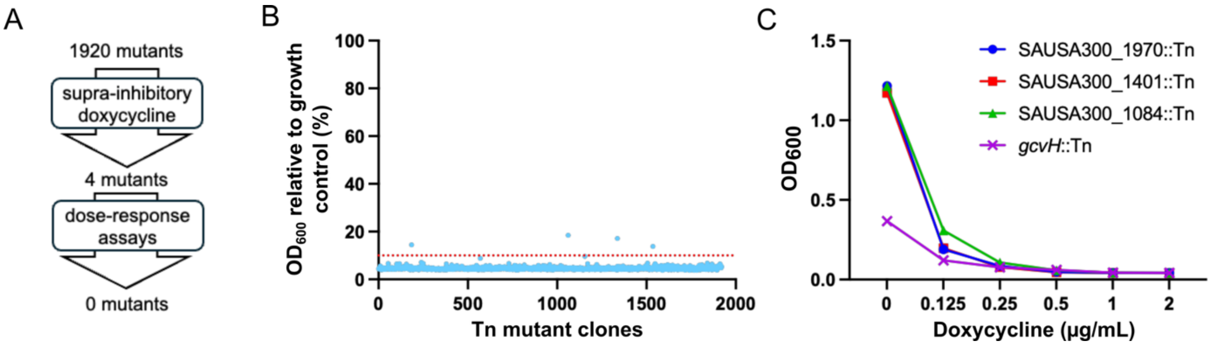


**Fig. S1. | Chemogenomic screen to identify susceptibility determinants of doxycycline in MHB.**

A) The workflow of the chemogenomic screen and follow-up assays that led to no putative determinants of doxycycline resistance in MHB. B) A chemogenomic screen of the Nebraska Transposon Mutant Library with doxycycline at 1 µg/mL (8X MIC) in MHB. C) Dose-response assays of putative determinants uncovered through (B).

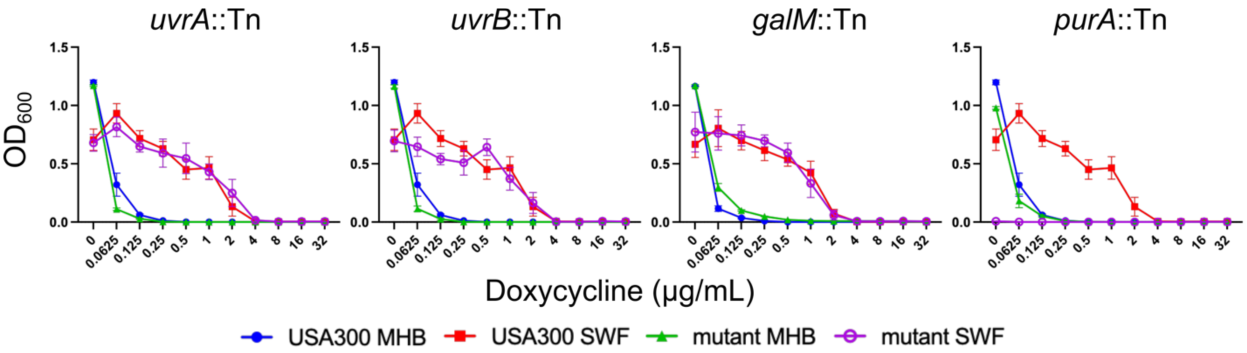


**Fig. S2 | MICs of mutants identified from the sub-inhibitory doxycycline screen that did not show increased susceptibility to doxycycline in SWF.**

The sub-inhibitory doxycycline screen is shown in Fig. 1. n = 4 from two independent experiments, shown as mean ± SEM.

**galM*::Tn inoculum adjusted to 0.01OD to improve growth in SWF growth control. *purA*::Tn did not grow in SWF, suggesting *purA* is conditionally essential in SWF.

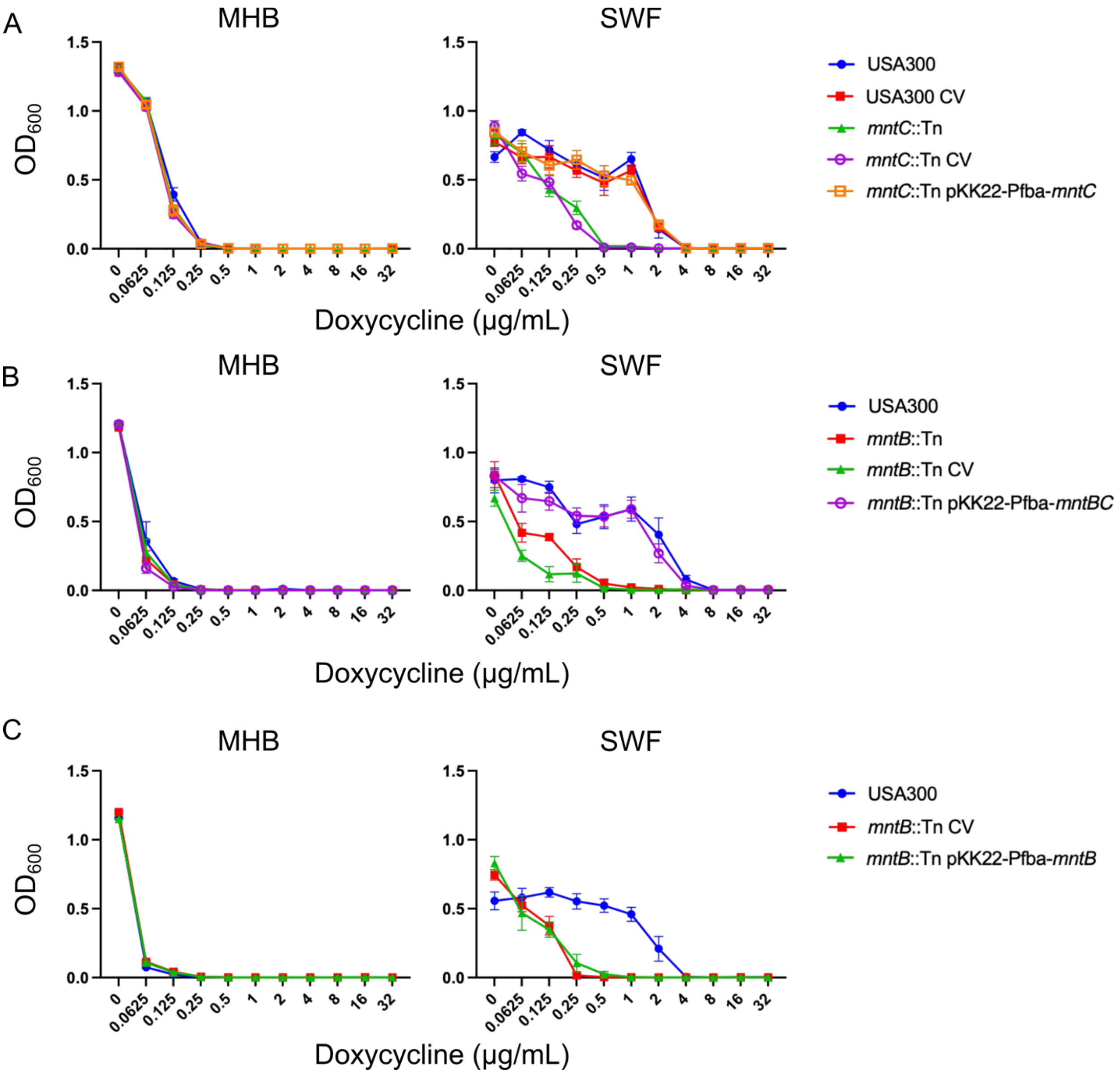


**Fig. S3 | Genetic complementation of *mntB*::Tn and *mntC*::Tn.**

A) MIC of doxycycline against USA300 and *mntC*::Tn with no vector, control vector (CV, parent vector lacking gene insert), or genetic complementation of *mntC*::Tn in both MHB and SWF. B) MIC of doxycycline against USA300 and *mntB*::Tn with no vector, CV, or genetic complementation of *mntB*::Tn with *mntBC* tested in both MHB and SWF. A, B) n = 6 from three independent experiments, shown as mean ± SEM. C) MIC of doxycycline against USA300 and *mntB*::Tn with CV, or genetic complementation of *mntB*. n = 4 from two independent experiments, shown as mean ± SEM.

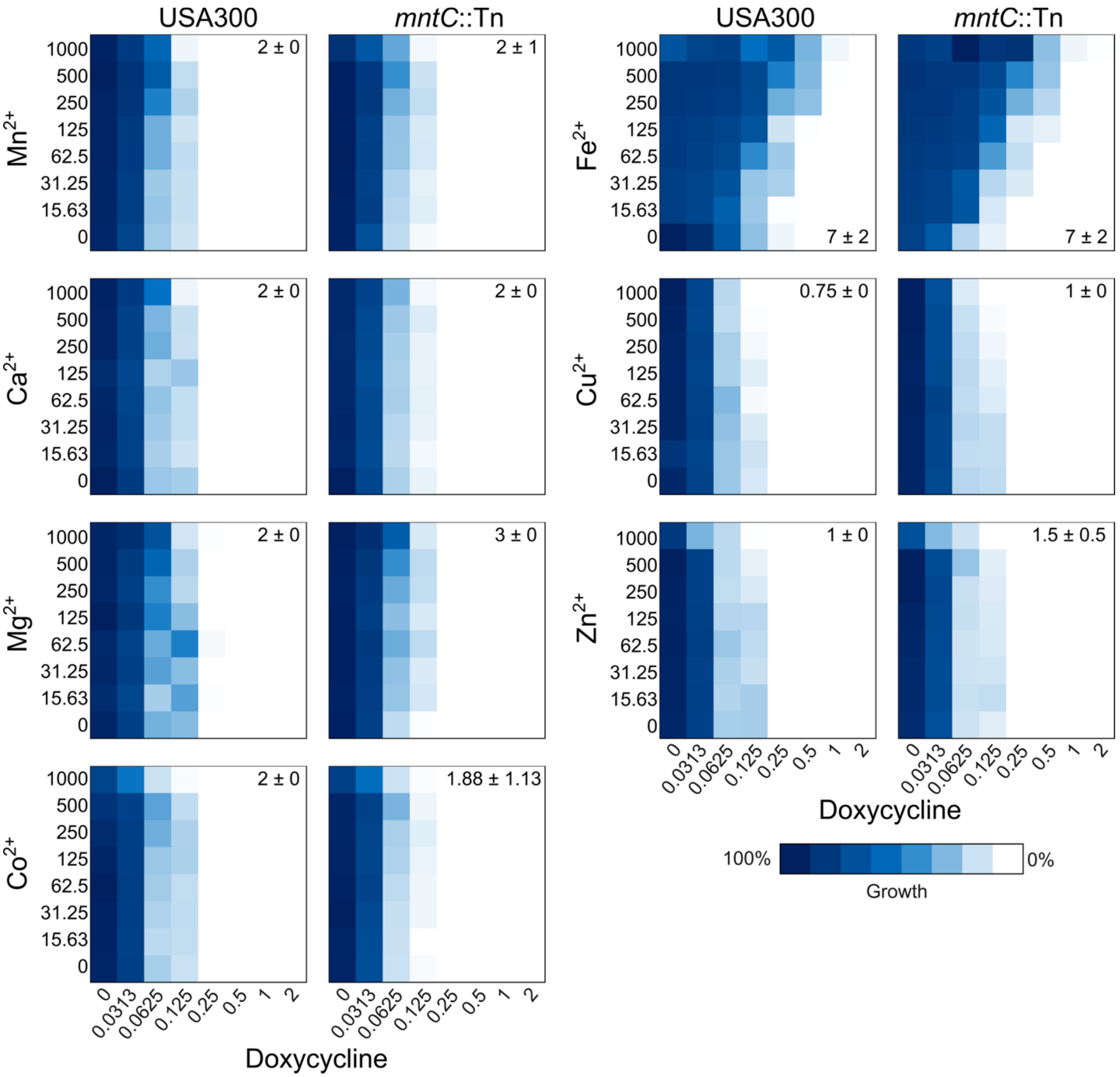


**Fig. S4 | Checkerboard assays of divalent cations with doxycycline.** Representative checkerboard assays of manganese, calcium, magnesium, cobalt, iron, copper, and zinc along the y-axis in µM and doxycycline (x-axis) in µg/mL against USA300 and *mntC*::Tn in MHB. n = 2, FICI values are displayed at the corner of each heatmap shown as mean ± SEM.

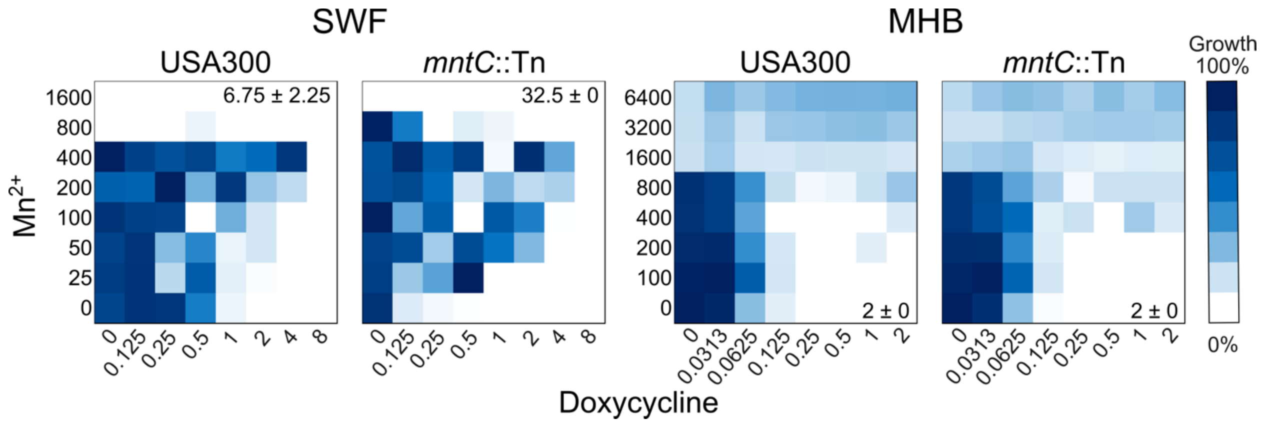


**Fig. S5 | Checkerboard assays of $Mn^{2+}$ with doxycycline.**

Representative checkerboard assays of manganese in µM and doxycycline in µg/mL against USA300 and *mntC*::Tn in MHB and SWF. n = 2, FICI values are displayed at the corner of each heatmap shown as mean ± SEM.

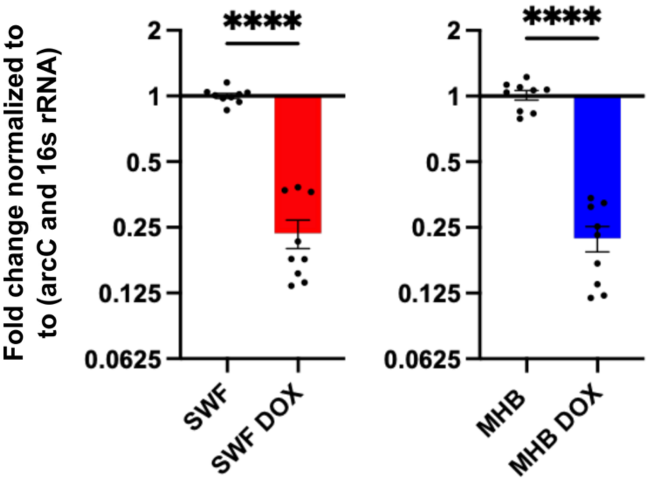


**Fig. S6 | Alteration in *mntC* transcription with and without doxycycline exposure.**

Fold change in *mntC* transcription from USA300 in SWF or MHB with or without doxycycline. n = 9 from three independent experiments. Significant differences were determined between USA300 cultured in MHB or SWF with and without doxycycline by Welch's t-test. p <0.0001 (****), p <0.001 (***), p <0.01 (**), p <0.05 (*).

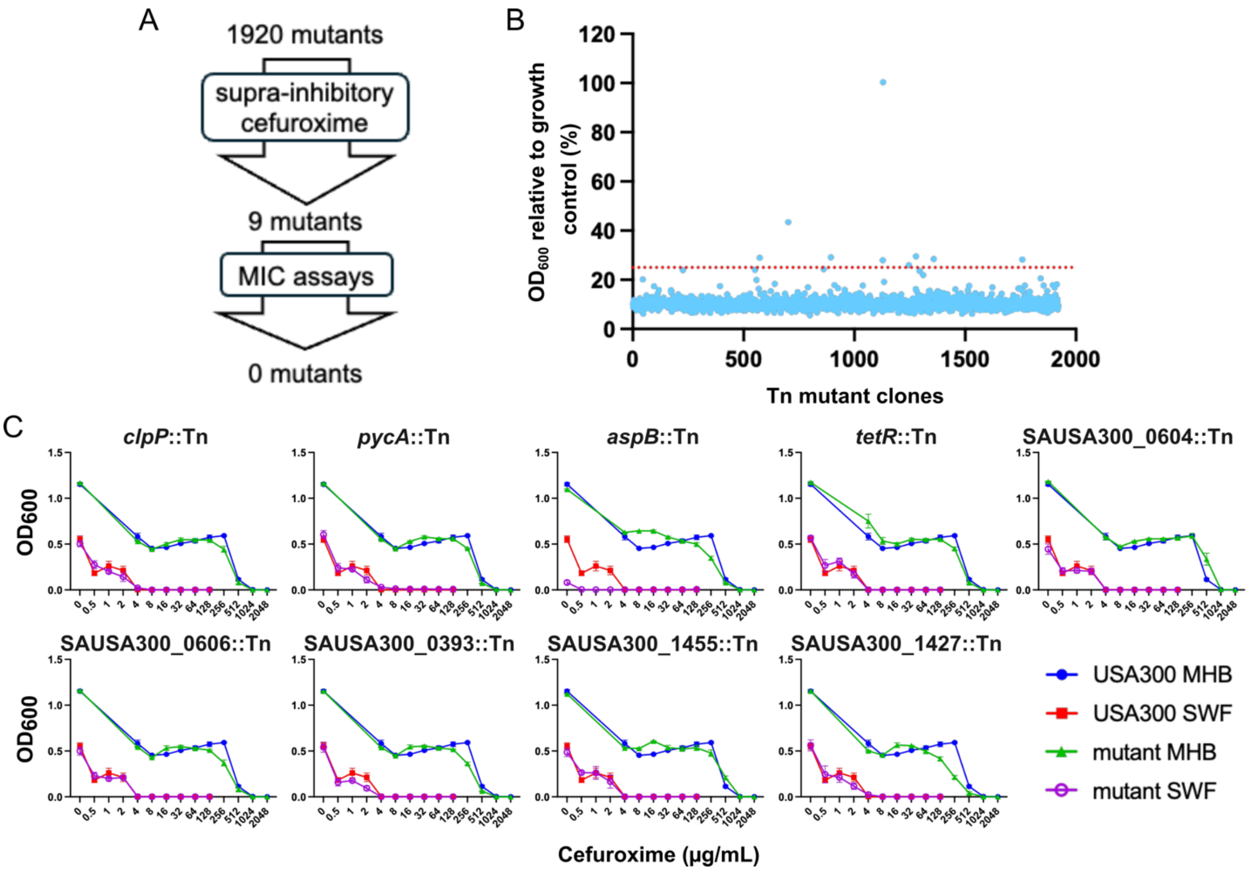


**Fig. S7 | Chemogenomic screen to identify susceptibility determinants of cefuroxime in SWF.**

A) The workflow of the chemogenomic screen and follow-up assays that led to no putative determinants of cefuroxime susceptibility in MHB. B) A chemogenomic screen of the Nebraska Transposon Mutant Library with cefuroxime at 16 µg/mL (4X MIC) in SWF. C) MIC assays of putative determinants uncovered through (B). n = 4 from two independent experiments, shown as mean ± SEM.

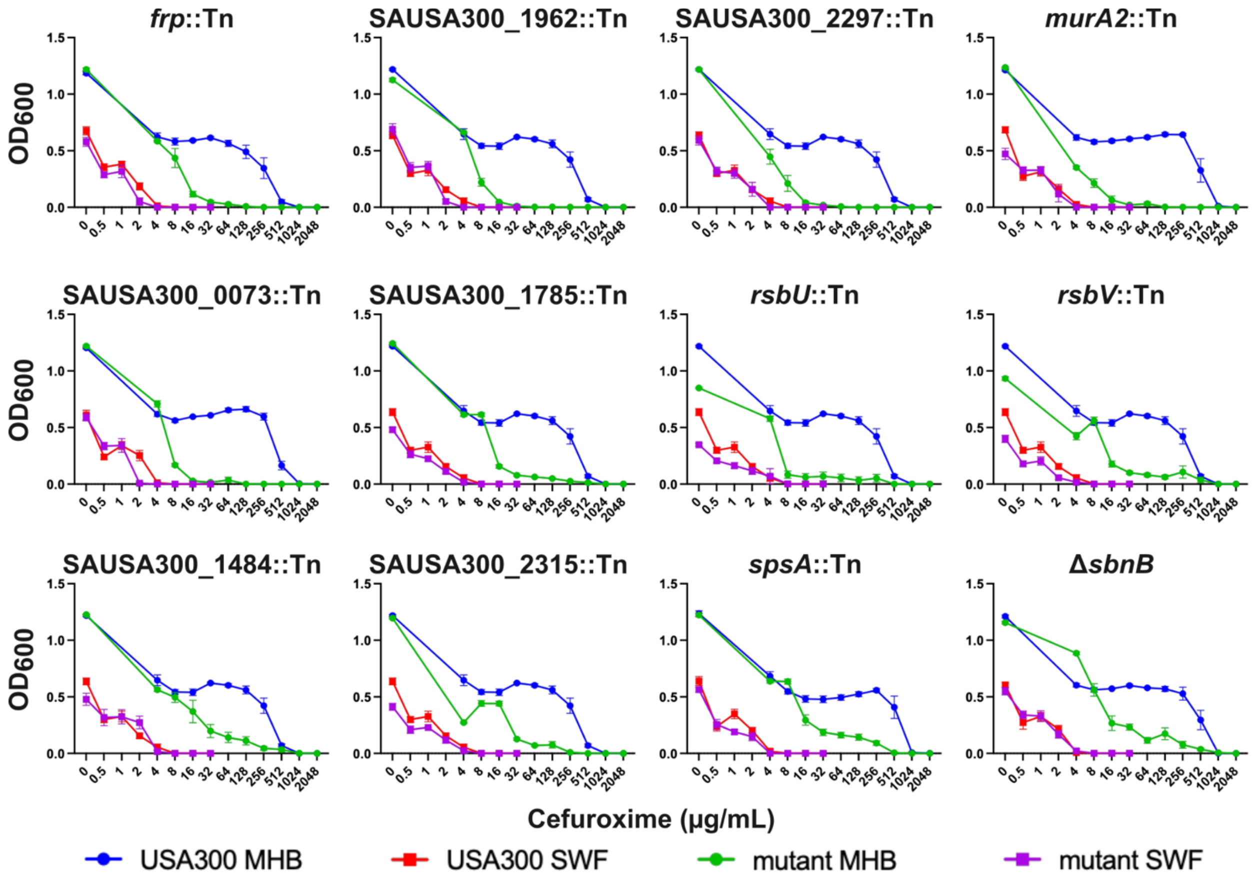


**Fig. S8 | MICs of putative secondary cefuroxime sensitivity determinants in SWF.** MIC assays of mutants, identified from the sub-inhibitory cefuroxime screen (Fig. 2), that showed a shift in cefuroxime MIC in MHB (that was more modest relative to that of mutants shown in Fig. 2C), potentially identifying secondary determinants of the phenotype. n ≥ 6 from at least three independent experiments shown as mean ± SEM.

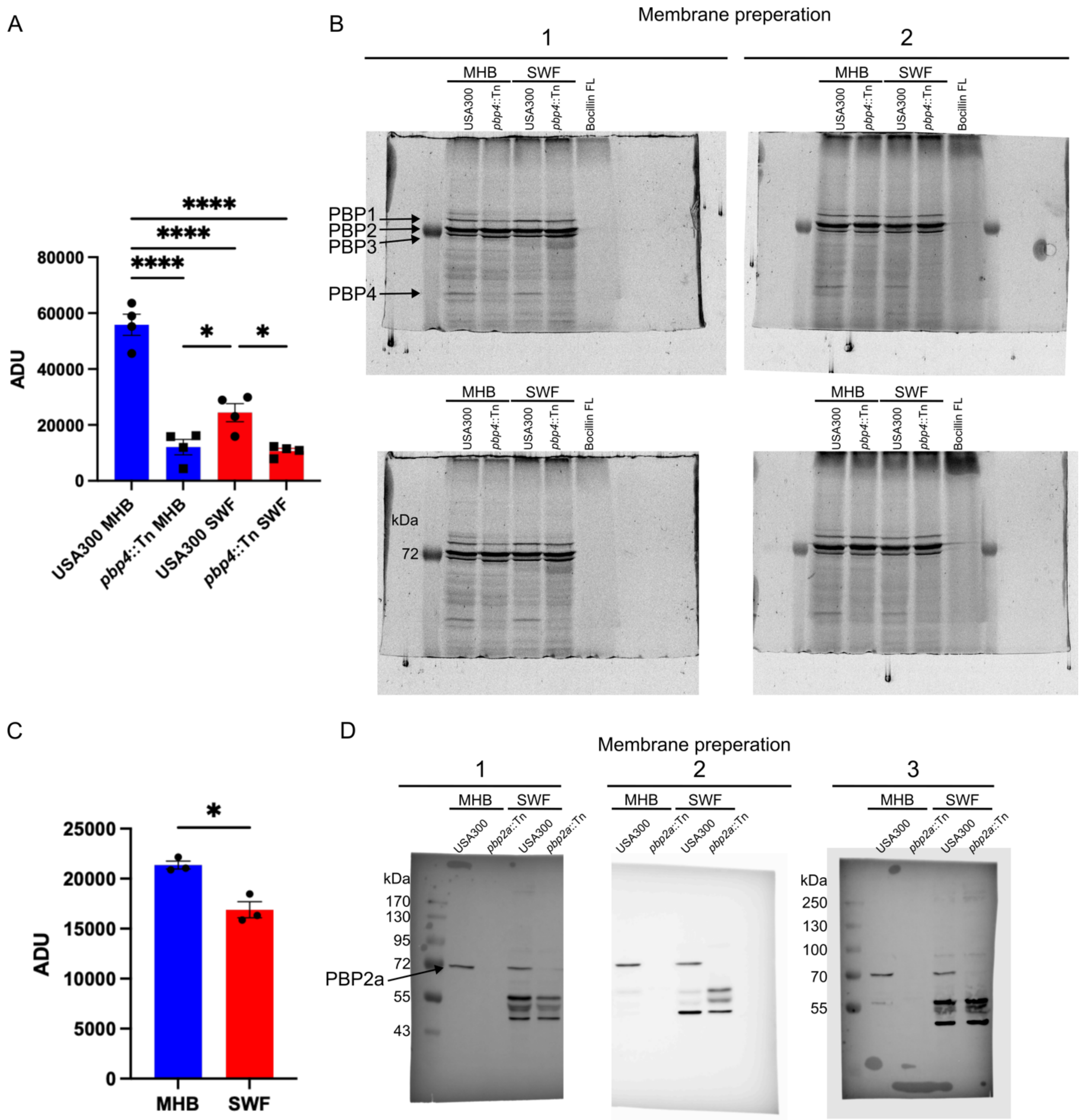


**Fig. S9 | Quantification of PBP1, 2, 3, 4, and PBP2a.**

A) PBP4 quantification displayed in arbitrary density units (ADU). Data from four gels from two independent membrane isolations. B) Uncropped gels of USA300 cell membranes stained with BOCILLIN FL isolated from: USA300 cultured in MHB, *pbp4*::Tn cultured in MHB, USA300 cultured in SWF, and *pbp4*::Tn cultured in SWF, in addition to a negative control of BOCILLIN FL with no membrane. C) PBP2a quantification displayed in ADU from three membrane isolations. D) Western blots, using an anti-PBP2a antibody, of membranes isolated from: USA300 cultured in MHB, *pbp2a*::Tn cultured in MHB, USA300 cultured in SWF, and *pbp2a*::Tn cultured in SWF. Significant differences were determined between wild-type and mutants cultured in MHB and SWF by two-way ANOVA and the Tukey post hoc test (in A), and by Welch's t-test (in C). $p < 0.0001$ (****), $p < 0.001$ (***), $p < 0.01$ (**), $p < 0.05$ (*).

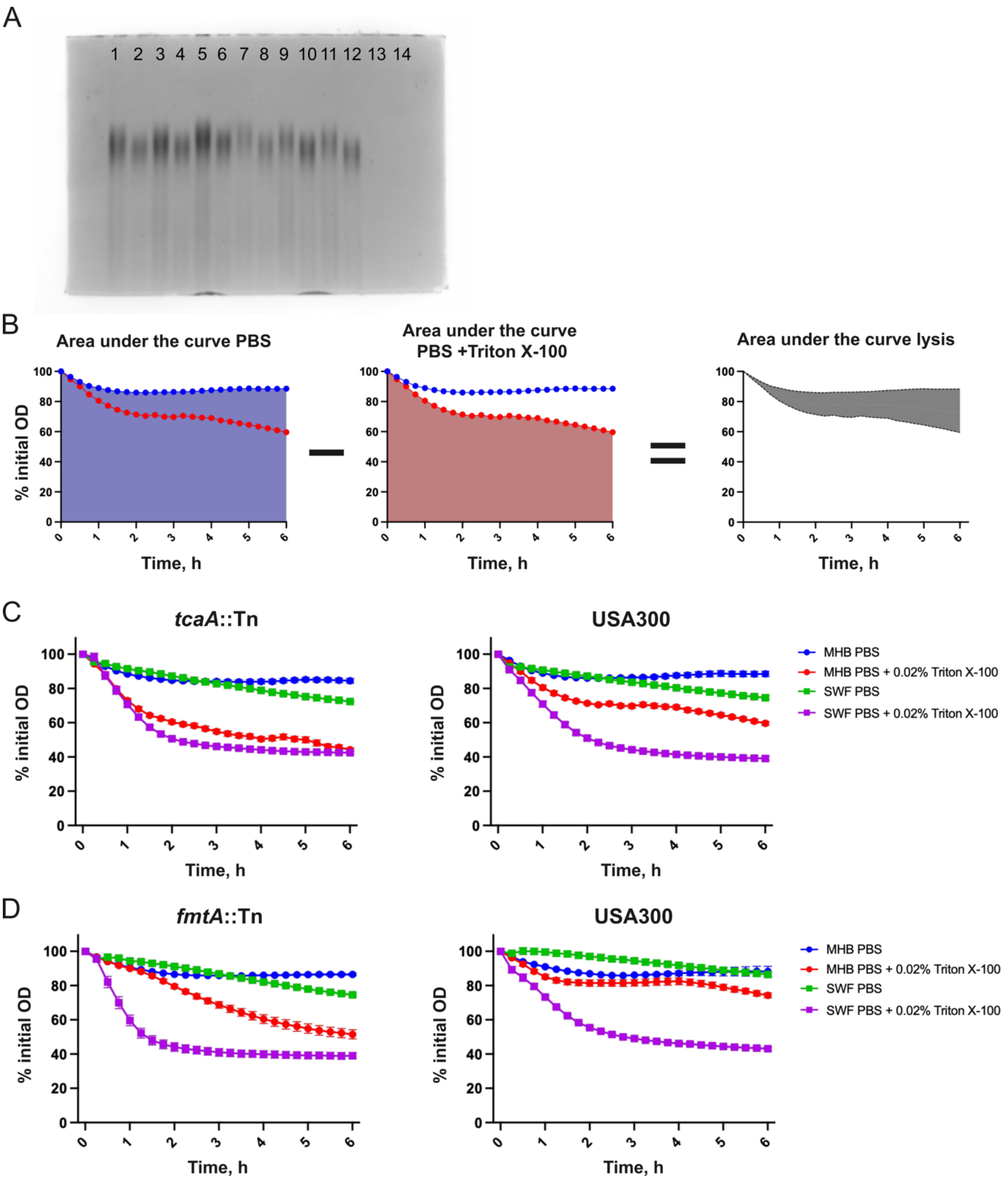


**Fig. S10 | WTA visualization and lysis curves of wild-type USA300 and the *tcaA*::Tn and *fmtA*::Tn mutants.**

A) Uncropped gel of WTA isolated from USA300 cultured in MHB (Lanes 1, 3, 5), USA300 cultured in SWF (Lanes 2, 4, 6), *tcaA*::Tn cultured in MHB (Lanes 7, 9, 11), and *tcaA*::Tn cultured in SWF (Lanes 8, 10, 12). B) Diagram illustrating how the area under the curves for lysis rates were calculated. C) The rate of lysis of *tcaA*::Tn and USA300 in PBS or PBS + 0.02% Triton X-100 cultured from MHB or SWF. n = 12 from four independent experiments, shown as mean ± SEM. D) The rate of lysis of *fmtA*::Tn and USA300 in PBS or PBS + 0.02% Triton X-100 cultured from MHB or SWF. n = 6 from two independent experiments, shown as mean ± SEM.

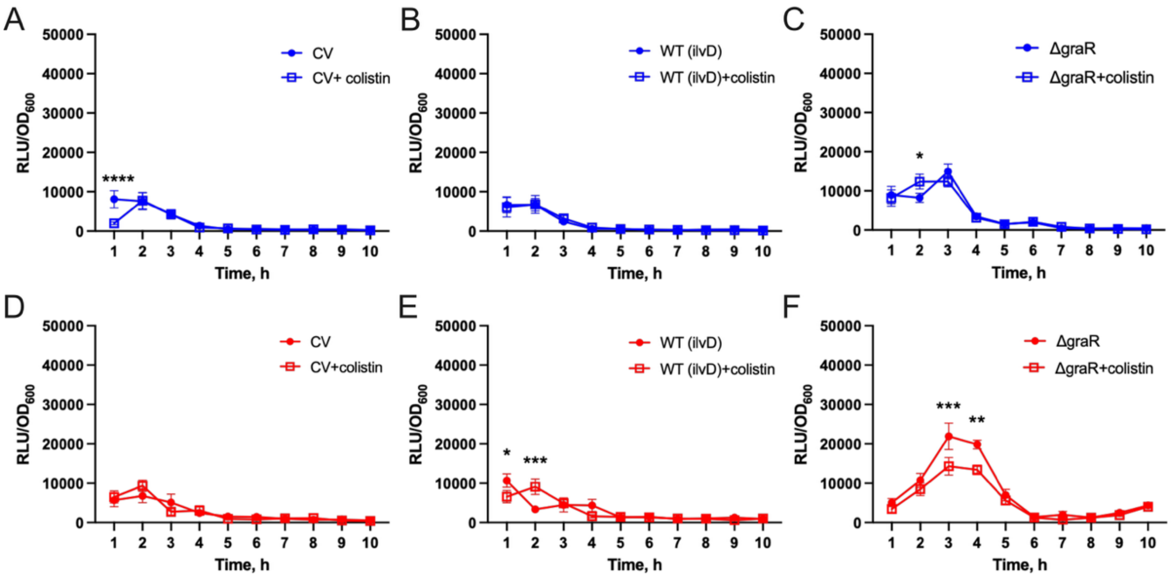


**Fig. S11 | Promoter-luciferase reporter activity of control strains for Fig. 4A.**
Monitoring the expression of luciferase from a promoterless control vector (CV) in USA300 (A, D), under the control of the *ilvD* promoter in USA300 (B, E), and under the control of the mprF promoter in Δ*graR* (C, F), monitored over 10 h in TSB (Blue) or SWF (Red) in the presence or absence of 1/32 MIC of colistin. n = 4 shown as mean ± SEM. Significant differences between strains treated with and without colistin were determined by two-way ANOVA and the Bonferroni post hoc test. p <0.0001 (****), p <0.001 (***), p <0.01 (**), p <0.05 (*).

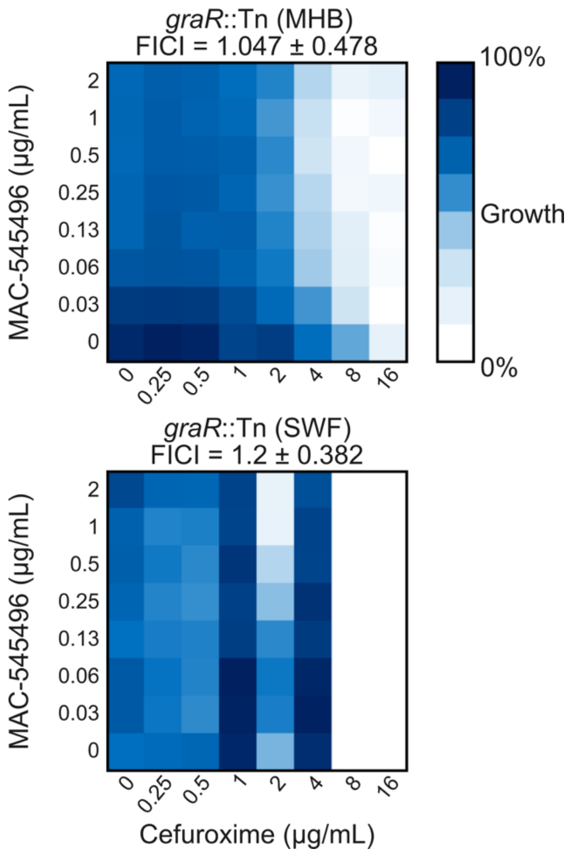


**Fig. S12 | Checkerboard assays of MAC-545496 and cefuroxime**

Representative checkerboard assays of *S. aureus graR*::Tn of cefuroxime in combination with MAC-545496. Dark blue represents high cell density measured by $OD_{600}$; n = 3; FICI shown as mean ± SEM.

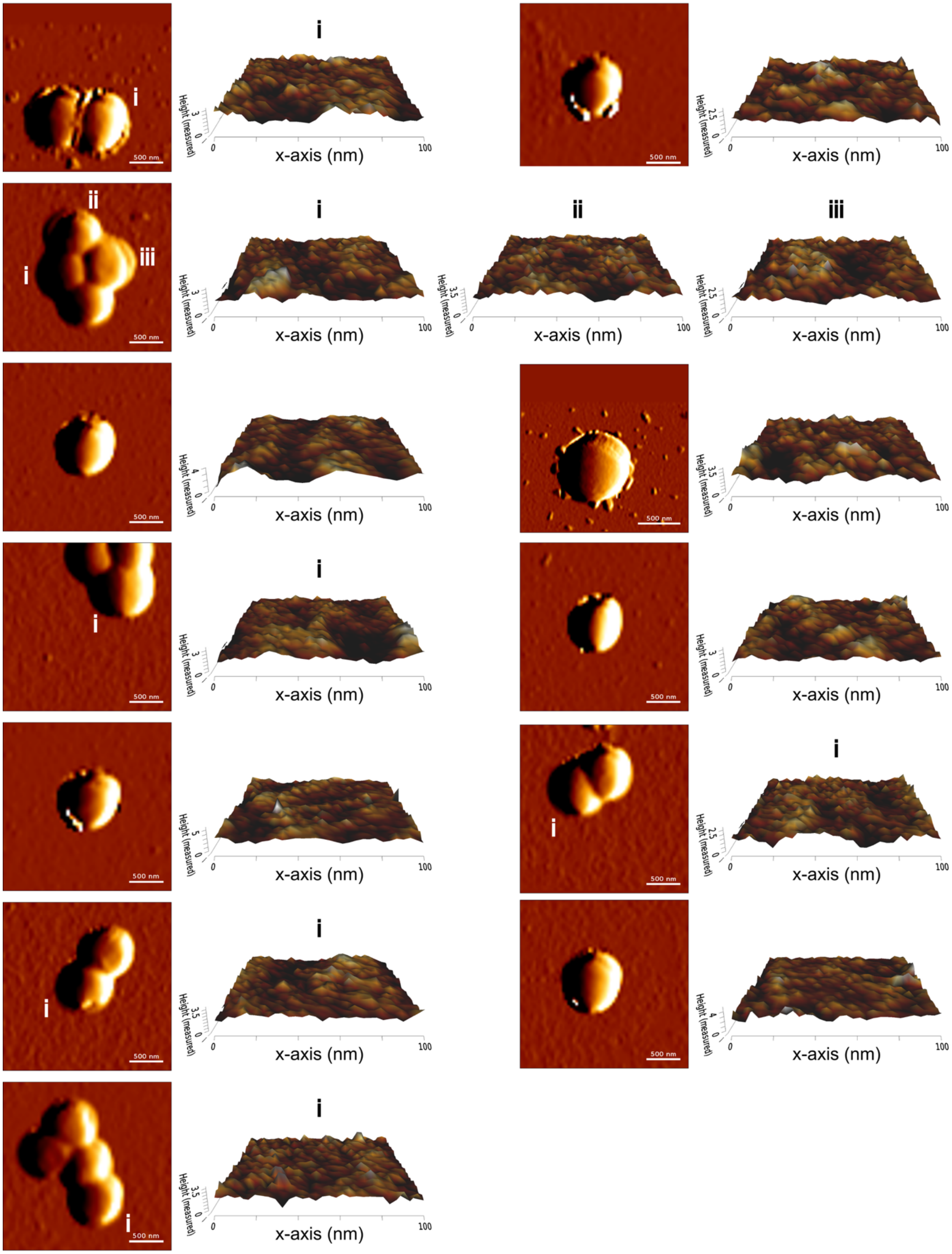


**Fig. S13 | AFM images of USA300 cultured in MHB**

Shown are 2.5 µm x 2.5 µm images of USA300 cultured in MHB and corresponding 100 nm x 100 nm 3D projections of the flattest section from the center of each cell.

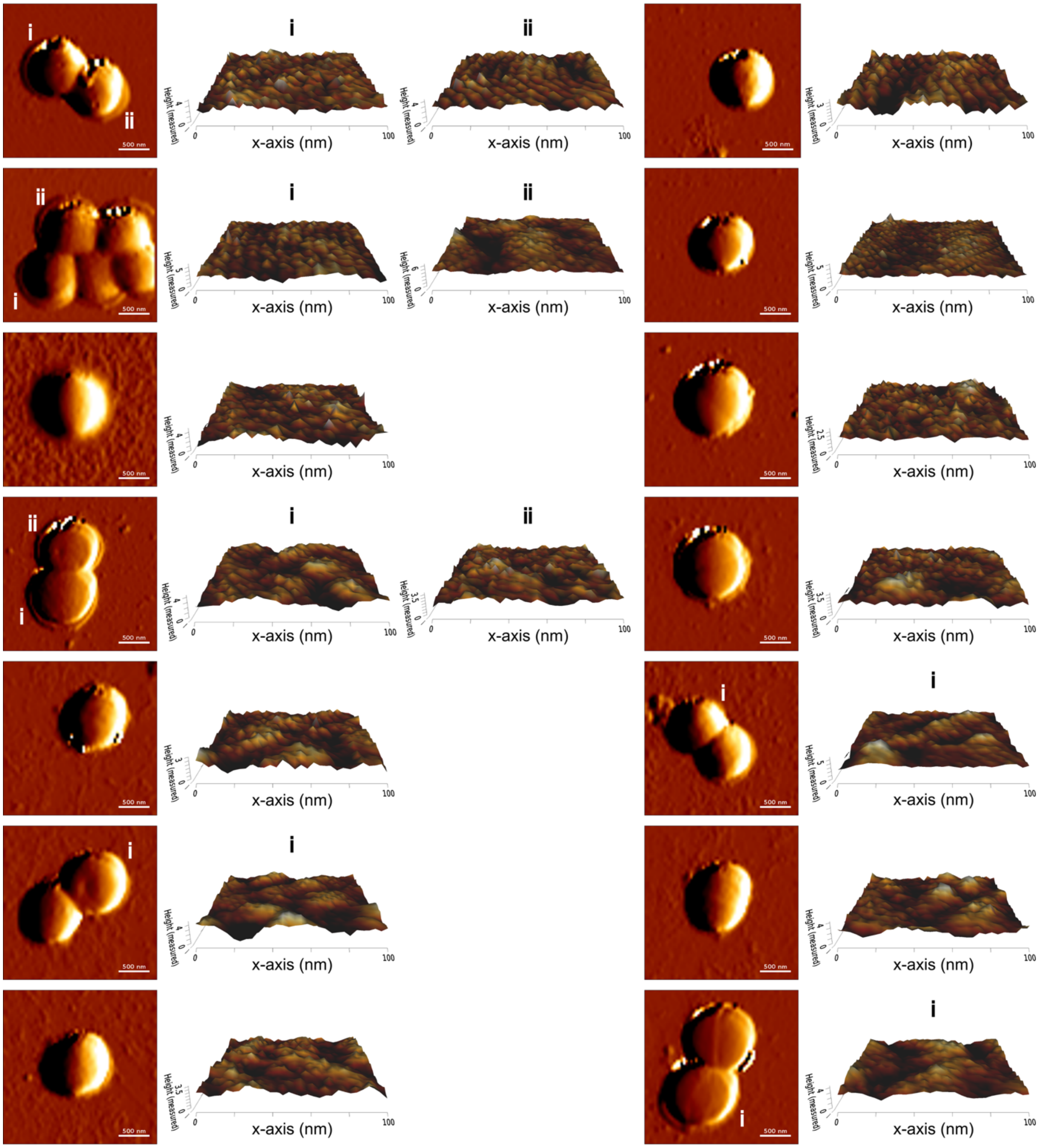


**Fig. S14 | AFM images of USA300 cultured in SWF**

Shown are 2.5 µm x 2.5 µm images of USA300 cultured in SWF and corresponding 100 nm x 100 nm 3D projections of the flattest section from the center of each cell.
